# Internet images reveal bumblebee-mimicking hoverflies have evolved to follow flower colour preferences of their models

**DOI:** 10.1101/2025.02.03.634701

**Authors:** James D. J. Gilbert, Joseph Craig-Jackson, Lesley Morrell, Francis Gilbert

## Abstract

Batesian mimics, such as hoverflies, can gain protection by resembling noxious species that predators avoid, such as bees. Such resemblance can be behavioural as well as morphological. Bees commonly have a preference for blue flowers, whilst flies generally prefer yellow and white. We predicted that, driven by selection for enhanced mimicry, bee-mimicking hoverflies would have switched flower preference from yellow to blue, to follow the flower choices of their models. We gathered internet photographs of flower-visiting bees and wasps, their hoverfly mimics, and non-mimicking hoverflies, and compared the colours of the visited flowers. Flowers visited by bumblebees were “bluer” than those visited by other hymenopterans (honeybees, wasps and solitary bees). Correspondingly, flowers visited by bumblebee-mimicking hoverflies were bluer than those visited by non-mimics, and were indistinguishable in blueness from those visited by bumblebees. There was no such pattern in flower “yellowness”, where all flies and most hymenopterans preferred similar yellowness, and only bumblebees chose less yellow flowers than the others. Our study demonstrates widespread changes in microhabitat choice (specifically flower preference) among a diverse group of Batesian mimics, consistent with the idea of behavioural mimicry. Mildly noxious models such as bumblebees are thought to exert particularly strong selection for mimetic accuracy, suggesting that habitat-choice mimicry may be selected as a way of enhancing morphological mimicry.

## Introduction

Batesian mimics such as hoverflies (Diptera:Syrphidae) are palatable to predators, but gain protection by resembling noxious model species that predators avoid (Rettenmeyer, 1970), such as bees (Hymenoptera:Apoidea). Mimics often experience strong directional selection to maintain or increase similarity to models (Mappes & Alatalo, 1997; Sherratt, 2002). To do this they mimic cues that predators normally use to recognise noxious prey, which can be subtle (Ruxton, 2009). For example, many species employ behavioural as well as morphological mimicry (Ceccarelli, 2008; Kitamura & Imafuku, 2015).

Bees and flies are both pollinators, and are typically associated with distinctive suites of floral characteristics (”pollination syndromes”), including distinct floral colours (Faegri & van der Pijl, 1979). Bees have long been known to prefer blue flowers (Giurfa et al., 1995; Gumbert, 2000; Lubbock, 1882; Raine et al., 2006; Raine & Chittka, 2007; Robertson, 1928), while flies (Diptera) prefer visiting yellow and/or white flowers (de Buck, 1990; Lunau, 2014).

We suggest that selection on mimetic accuracy may include selection for mimetic flies to mimic the flower choices of their models. A large part of the predation of dipteran species takes place on and around flowers (Dlusski, 1984), and closer mimics are attacked by predators at a higher frequency (F. Gilbert, 2005; Morse, 1979). Association with “fly flowers” could potentially be used by predators as a cue to a mimic’s identity. As one potential example, bee-flies (Bombyliidae) are known for their exceptionally close mimicry of bumblebees, and unusually for flies, they also prefer visiting blue and violet flowers (Kastinger & Weber, 2001); so much so, in fact, that they have evolved long tongues with which to exploit them (Szucsich & Krenn, 2000).

Hoverflies are an ideal clade in which to test this hypothesis. The Syrphidae contain several origins of mimicry of various hymenopterans (Leavey et al., 2021), varying from imperfect to astonishingly accurate (Gilbert 2005). While hoverflies commonly resemble their models morphologically, in some species elaborate behavioural mimicry has also been documented (Golding & Ennos, 2005) such as mock stinging, leg waving and wing wagging (Penney et al., 2014). In many cases floral visitation patterns of syrphid mimics follows that of their models, whether seasonally (Howarth & Edmunds, 2000), daily (Howarth et al., 2004) or in second-to-second foraging patterns (Golding & Edmunds, 2000).

Like other flies, syrphids are well known to have a general and presumably ancestral preference for yellow and white flowers (Hannah et al., 2019; Ilse, 1949; Kugler, 1950; Lunau et al., 2018; Robertson, 1928; Rodríguez-Gasol et al., 2019), and only rarely visit blue flowers (Ssymank, 1989, 2001). We predicted that hoverflies mimicking hymenopteran species that prefer blue flowers, such as bees, should have switched their own preferences towards bluer flowers than non-mimics, or related species that mimic other models. In addition, behavioural mimicry has mostly evolved among larger and morphologically closer mimics (Penney et al., 2014), lending credence to the established idea that larger body size selects for more faithful mimicry (Penney et al., 2012). Thus, we also predicted that flower preferences are most likely to have switched in the largest hoverflies.

For assessing visual traits of organisms, images obtained from the internet have been shown to provide a good substitute for field data which can be difficult to obtain (Leighton et al., 2016). This can include studies assessing colour-based traits (Atsumi & Koizumi, 2017; Austen et al., 2018; M. P. Moore et al., 2019) despite variation due to camera- and environment-related differences (Laitly et al., 2021; Stevens et al., 2007). In this study, we obtained internet images of flower-visiting syrphid mimics and their hymenopteran models, as well as non-mimetic syrphids. We extracted data from those images on the RGB colour of the flowers being visited. We then used these data to test the hypotheses that (a) mimetic syrphids should follow the flower colour choices of their hymenopteran models, rather than the yellow/white preferences of their dipteran (and non-mimetic syrphid) ancestors, and (b) larger bodied syrphids should show greater fidelity to the colour choices of their models.

## Methods

Broadly speaking, we collected images depicting insects on flowers, and processed them to remove digital colour profiles (see below). Then, we digitally cut out the areas within the images showing the flowers, calculated their mean RGB colours and compared these values statistically among groups. By using internet-obtained images, we accepted that we had to ignore colours outside of the human-visible spectrum that might be relevant to predators, e.g. in the UV range; and that unknown camera types and settings result in a degree of uncontrolled colour bias in all photographs (Laitly et al., 2021; Stevens et al., 2007). We made the simplifying assumption that photographs would not be systematically biased when depicting different groups (such as models and mimics) towards particular camera settings or lighting conditions. For example, we assumed that models and mimics would not systematically be photographed at different times of day in a way that would affect the imaged flower colour.

### Image collection

Mimetic syrphid species were matched to putative Hymenopteran models (Table S1) using a near-exhaustive list of descriptions of their mimicry in syrphid-related literature, collated by FG. Each mimic species and its set of one or more putative model species were grouped as a “model-mimic pair” (e.g. *Eristalis pertinax* with *Apis mellifera*; *Volucella bombylans* var. *plumata* with a pooled set comprising *Bombus terrestris*, *B. lucorum*, *B. hortorum*, *B. soroensis* and *B. jonellus*; see Table S1).

Model-mimic pairs were then grouped into four distinct morphotypes, determined by the identity of the model identified in Table S1. These were: Bumblebee [Apoidea:*Bombus*], Honeybee [Apoidea:*Apis*], Solitary bee [all non-eusocial Apoidea i.e. not *Apis*, *Bombus* or other social bee], and Social wasp [Aculeata:Vespidae]). We also included a final morphotype “Non-mimic hoverflies” which were identified as such in the literature (e.g. *Ferdinandea cuprea*, *Baccha elongata*; Table S1).

Internet images showing flowers being visited by mimics, their corresponding models, and non-mimic hoverflies were obtained from web searches in March-April 2020, using common (e.g. “Tapered drone fly”) and binomial names (e.g. “Eristalis pertinax”) as search terms. For each of the four model morphotypes, we gathered photographs of 10 mimic and 10 model species with 10 photographs per species, as well as 5 non-mimic syrphid species with 20 photographs each. Thus, in total we gathered 100 photographs for each morphotype. For each morphotype we made sure to gather images of syrphids from across the phylogeny, and not from a single evolutionary origin (Leavey et al., 2021; see Figure 1).

**Figure 1.**
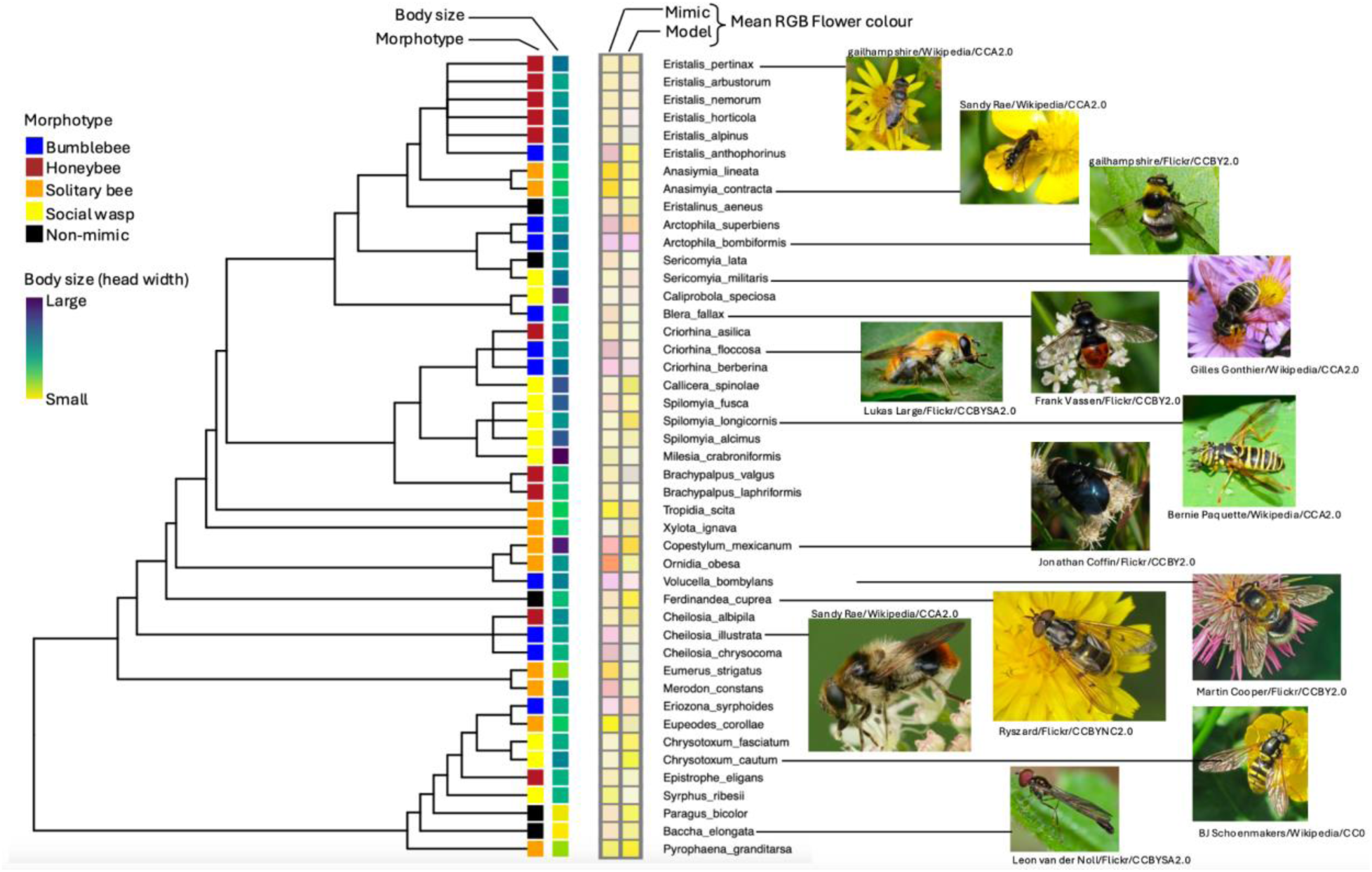
Syrphid phylogeny used in this study (with arbitrarily scaled branch lengths) showing morphotype, body size, and the flower colour preferences of mimics and models, as assessed by internet images depicting the species visiting a flower. Photos of representative species are given, but are not necessarily part of the image dataset.

All images were collected from nationally recognised nature ID and photography sites, such as bugguide (bugguide.net) and iNaturalist (inaturalist.org); specialised forums for ID of images of Diptera, such as Diptera.info (diptera.info); or sites with clear author attribution of images, e.g. Flickr (www.flickr.com) and where the author was a known authority, such as the Flickr gallery maintained by S. Falk (https://www.flickr.com/people/63075200@N07/). Images were discarded if the putative species ID could not be verified.

We chose only images showing the insect on a clearly visible flower and with dimensions above approximately 275x183 pixels to ensure sufficient number of pixels per image for effective flower colour comparison and consistency between monitors of varying resolution (Tkalcic and Tasic, 2003). While this limited the sample size in the case of some species with few available images of confirmed ID and sufficient quality, such as *Brachypalpus laphriformis,* this did not significantly affect the sample sizes per group (N=100 images) or, we assumed, average RGB values for each image.

### Image processing

We modified all images to reduce the effect of camera setting adjustments applied by camera profiles (exposure, brightness etc.) using the program RawTherapee (RawTherapee Team, 2020). To do this, we modified the RGB values obtained from each image directly, using RawTherapee’s “processing profile” function to remove any profile. Typically this brightened each image, bringing R, G and B values of consistent blocks of colour closer to ‘true’ R, G and B values (Stevens et al., 2007; Tkalcic & Tasic, 2003).

We then digitally extracted the part of each image containing the flower(s) using the GNU Image Manipulation Program (GIMP) 2.10.14 (The GIMP Team, 2020), using GIMP’s “free select” tool. We used the histogram tool with a linear axis to extract RGB profiles for each cutout, and recorded the median RGB values. A linear rather than logarithmic axis was used for this process (Stevens et al., 2007), since it is best suited for use with photographs, particularly those taken in the field, whereas a logarithmic axis is more commonly used for images containing substantial areas of constant colour (Smith & Joost, 2012).

### Body size

We used head width as a proxy for body size using a dataset, compiled by FSG, that combined our previously published data (F. S. Gilbert, 1985) with subsequent unpublished measurements. Twenty species were directly measured in this dataset (Table S2). For the 20 remaining species not directly measured, we interpolated species-typical head width from intra-generic regressions (typically very tight; F. S. Gilbert, 1985), using congeners in FSG’s dataset, based on other structural measurements such as wing length and body length taken from the literature (Bot & Van de Meutter, 2023; Skevington et al., 2019; Thompson, 1991).

### Data analysis

All analyses were conducted in R 4.4.1 (R Core Team, 2023). We log-transformed the original three RGB values of the flower cutouts to conform with linear assumptions. We then reduced their dimensions to scores along two principal axes of variation using Principal Component Analysis (Figure 2). Broadly speaking, PCA1 equated to yellow (a combination of red and green) and PCA2 to blue, and we refer to them henceforth as “yellow” and “blue” respectively.

**Figure 2.**
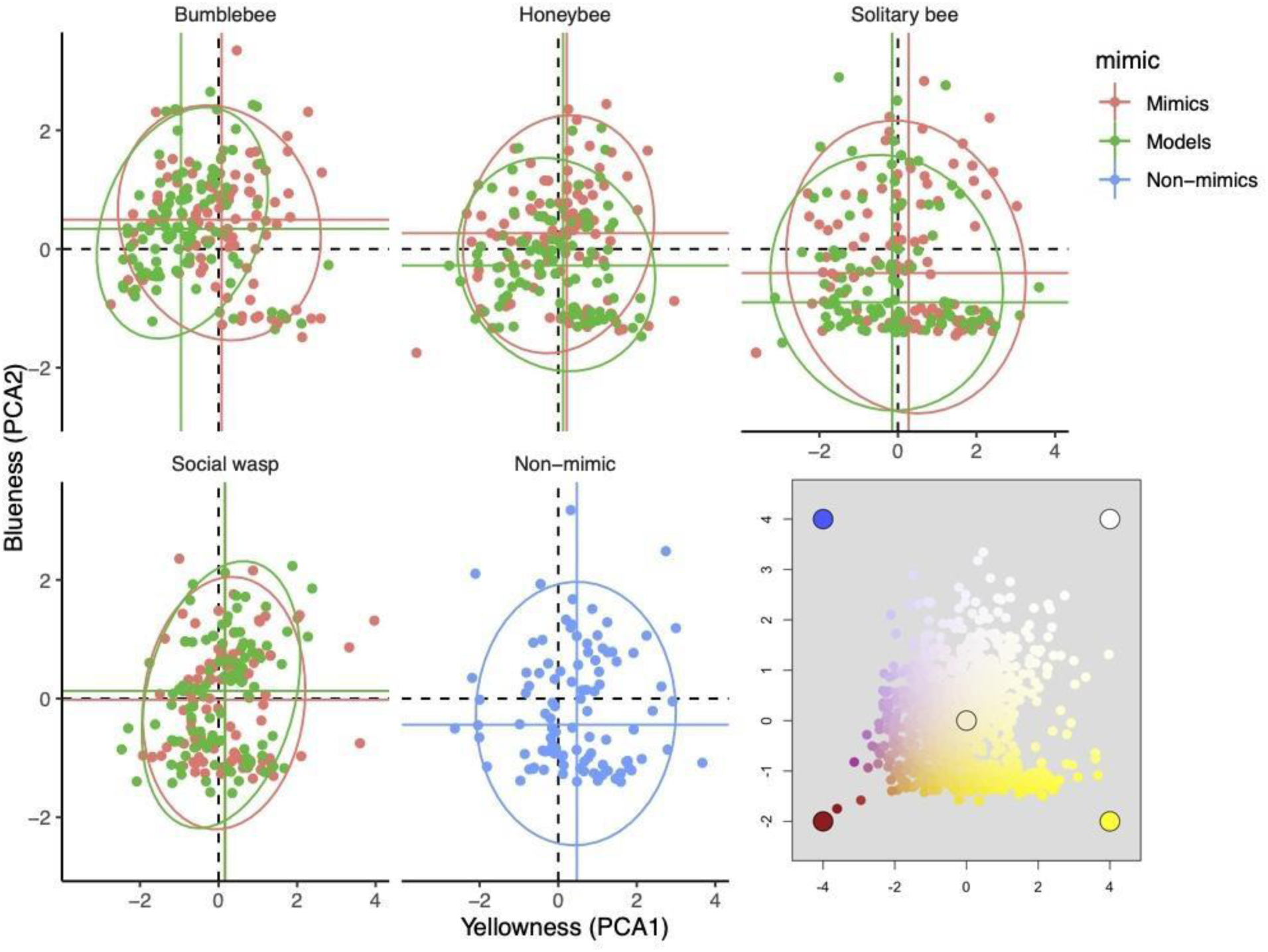
Blueness (PCA2) plotted against yellowness (PCA1) for models and mimics of different morphotypes, with means and 95% confidence ellipses shown in solid colours. Inset shows RGB colours for the whole dataset, with representative extremes marked for comparison.

To compare colour preferences, we first compared all syrphid morphotypes including non-mimics (bumblebee, honeybee, solitary bee, social wasp, and non-mimic) with model morphotypes, in a one-way analysis. We did not include phylogenetic information here because, when hymenopteran models were included, the evolutionary distances between syrphids and their models dwarfed the inter-species distances within these clades. We used mixed-effects models, implemented in the lme4 package in R (Bates, 2010), with either PCA1 or PCA2 as a response, “morphotype” as a fixed effect and “species” as a random effect. We used the emmeans package (Lenth, 2023) to conduct specific comparisons among morphotypes.

Then we looked specifically at the effect of “being a model versus a mimic” within the different morphotypes. For this, we trimmed the dataset to exclude non-mimic hoverflies, and fitted a mixed effect model with “morphotype” and “model or mimic” plus their interaction as fixed effects and “model-mimic species pair” as a random effect.

Finally, we investigated whether mimics had departed from non-mimics in their flower colour preference, and whether body size could explain any such departure. For this, we compared just the hoverfly species across the syrphid phylogeny, excluding hymenopteran models. We used the MCMCglmm package in R to fit the model while also accounting for nonindependence due to shared evolutionary history (the syrphid phylogeny), and also multiple observations for each tip of the phylogeny (by including species as a random effect in the model). We used “morphotype”, “body size” and their interaction as fixed predictors, with 100000 MCMC iterations, a thinning interval of 50 and a burnin of 2000, checking for convergence using trace plots and Gelman-Rubin diagnostics over 3 replicate chains. We used weakly informative priors with V=1 and nu=0.2 for both fixed and random effects.

The phylogeny (Figure 1) was developed following recommendations of Beaulieu et al (2012). The base phylogeny was based on Leavey et al (2021) at the genus level, with species within genera added as star phylogenies where necessary. Branch lengths for added species were unknown, so branch lengths across the tree were scaled arbitrarily according to node depth, following Grafen (1989), using a power of 0.8.

## Results

The 45 syrphids in the dataset, their mean colour preferences, and those of their models, can be seen in Figure 1. Flower colour data can be found in Table S3.

Two axes explained 83% of the variance in the colour of flowers photographed being visited by models or mimics. Axis 1 (49% of the variance) was aligned strongly and identically with red (71%) and green (70%) and will be known henceforth as yellowness. Axis 2 (34% of the variance) was exclusively aligned with blue (98%) and we refer to this axis as blueness.

Scores along the yellowness and blueness axes for different morphotypes (models and mimics) are shown in Figure 2.

Hymenopteran models of different morphotypes differed in the yellowness of the flowers they chose to visit, while syrphids (mimics and non-mimics) all chose flowers of similar yellowness (Linear mixed model, X2=32.5, df=8, p<0.0001, Figure 3). When we examined effects of mimicry and morphotype independently, excluding non-mimic hoverflies, there was a significant “mimic/model” x morphotype interaction (X2=8.45, df=3, p=0.037, Table 2). Bumblebees chose less yellow flowers than other models, which did not differ from each other (honeybee, solitary bee and social wasp). In contrast, all mimic syrphids chose a similar yellowness, which was also similar to that chosen by honeybees, solitary bees and social wasps, but was higher than the yellowness chosen by bumblebees.

**Figure 3:**
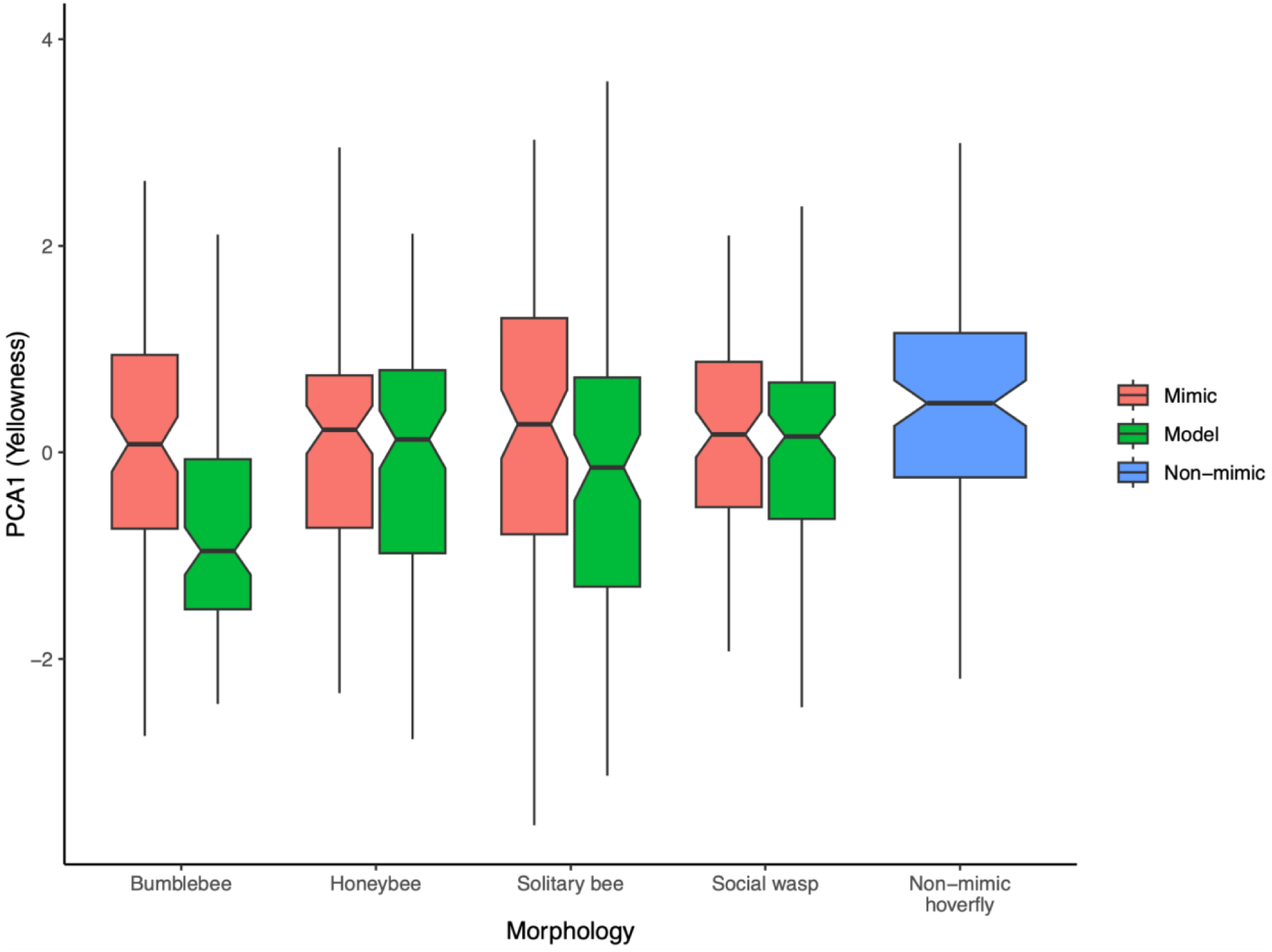
Yellowness preferences by morphotype and mimicry status. Syrphids (mimetic and non-mimetic) showed similar yellowness preference, while hymenopteran models differed (see text for details).

**Figure 4.**
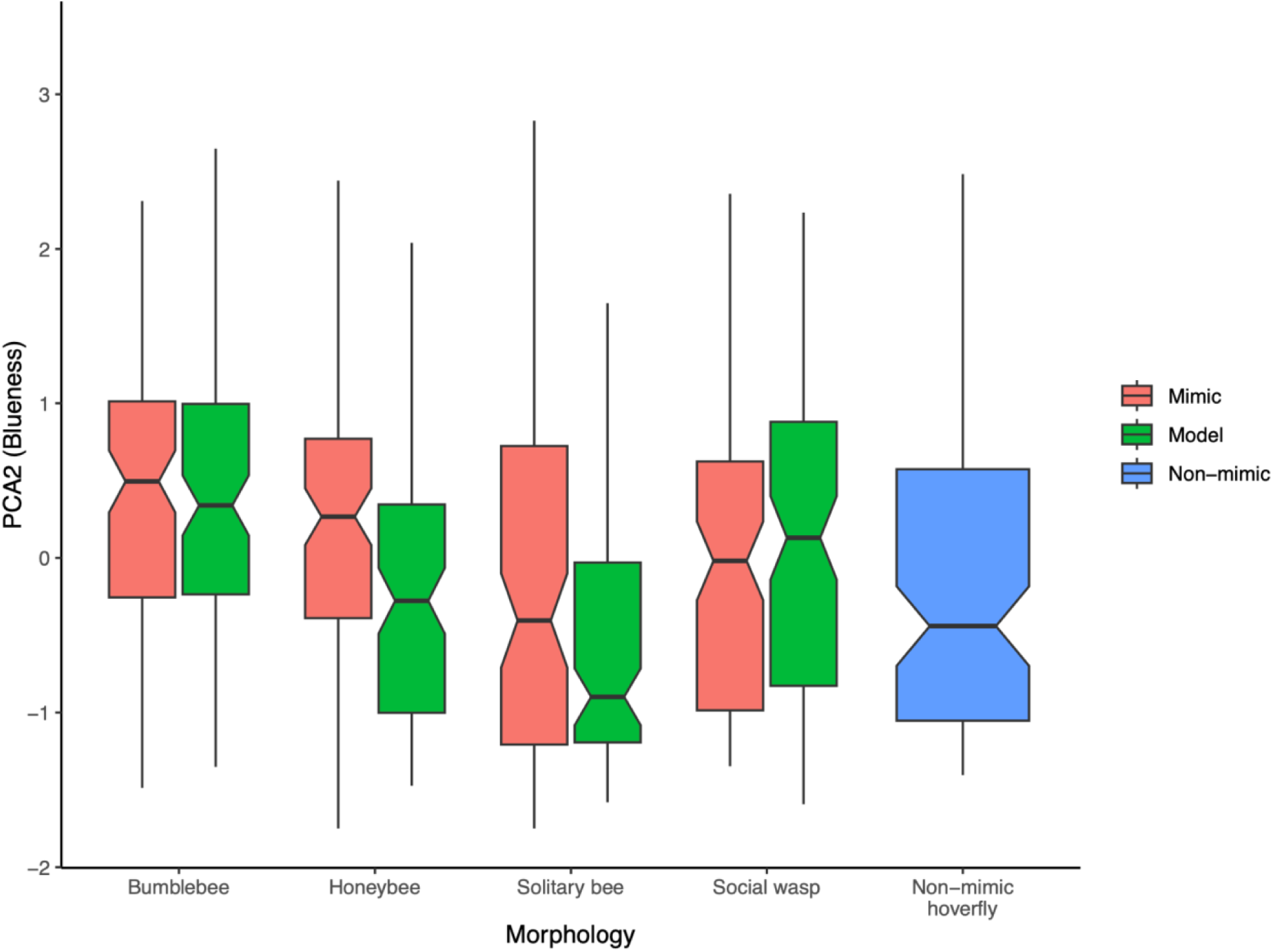
Blueness preferences by morphotype and mimicry status. Syrphid mimics were different from non-mimics, and tended to follow the preferences of their models such that morphotypes differed but models and mimics were statistically similar (see text for details).

**Table 1.**
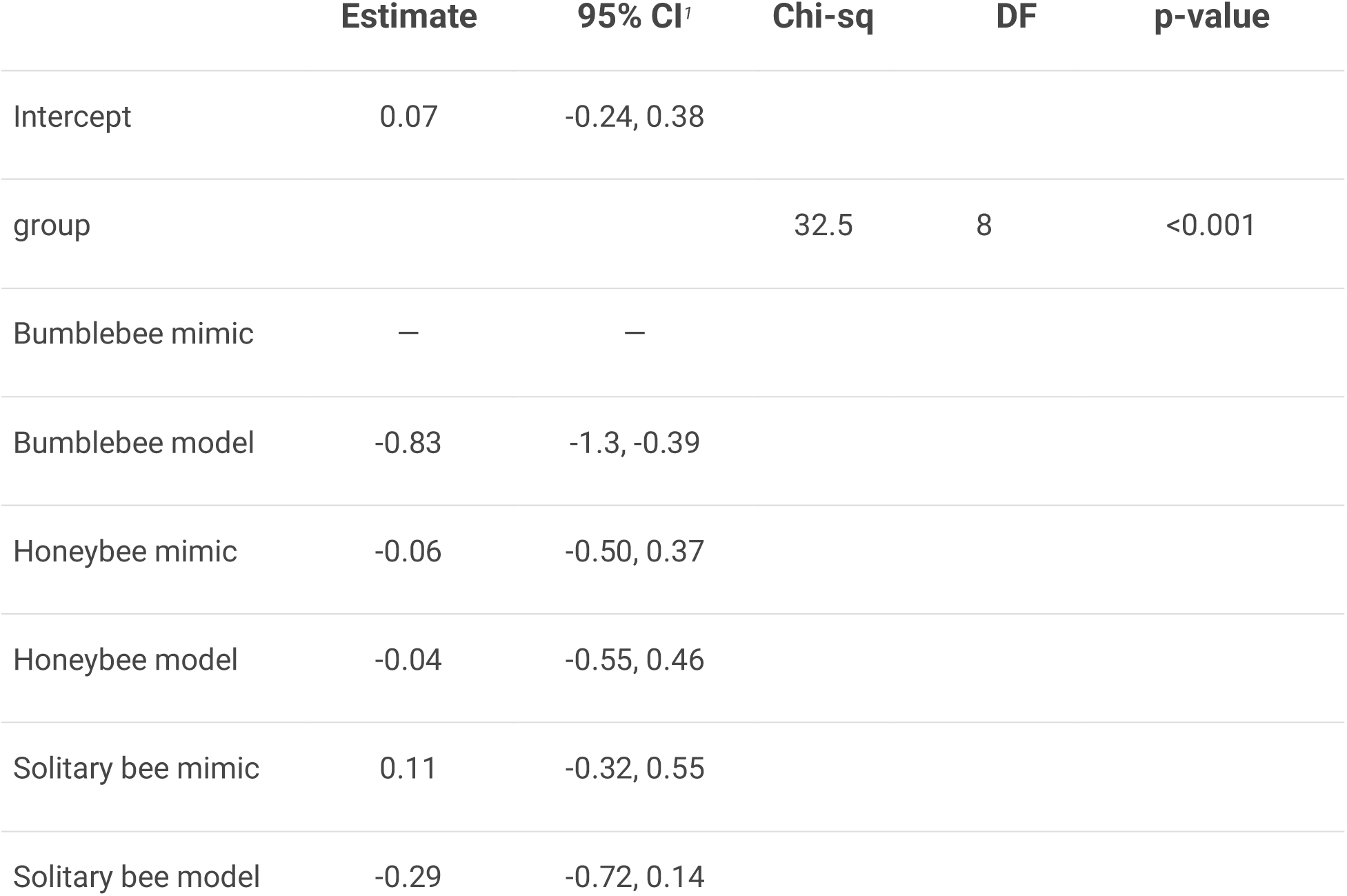
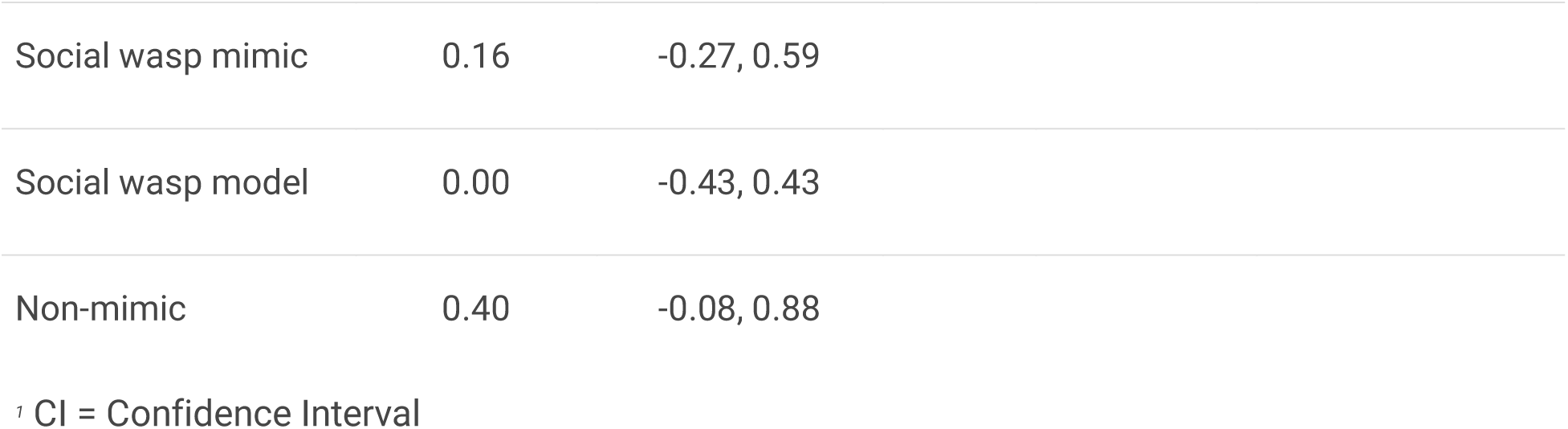
Linear mixed-effect model of flower yellowness against morphotype, with species as random effect.

|  | **Estimate** | **95% CI** | **Chi-sq** | **DF** | **p-value** |
| --- | --- | --- | --- | --- | --- |
| Intercept | 0.07 | -0.24, 0.38 |  |  |  |
| group |  |  | 32.5 | 8 | <0.001 |
| Bumblebee mimic | — | — |  |  |  |
| Bumblebee model | -0.83 | -1.3, -0.39 |  |  |  |
| Honeybee mimic | -0.06 | -0.50, 0.37 |  |  |  |
| Honeybee model | -0.04 | -0.55, 0.46 |  |  |  |
| Solitary bee mimic | 0.11 | -0.32, 0.55 |  |  |  |
| Solitary bee model | -0.29 | -0.72, 0.14 |  |  |  |

|  |  |  |
| --- | --- | --- |
| Social wasp mimic | 0.16 | -0.27, 0.59 |
| Social wasp model | 0.00 | -0.43, 0.43 |
| Non-mimic | 0.40 | -0.08, 0.88 |

<sup>1</sup> CI = Confidence Interval

**Table 2.**
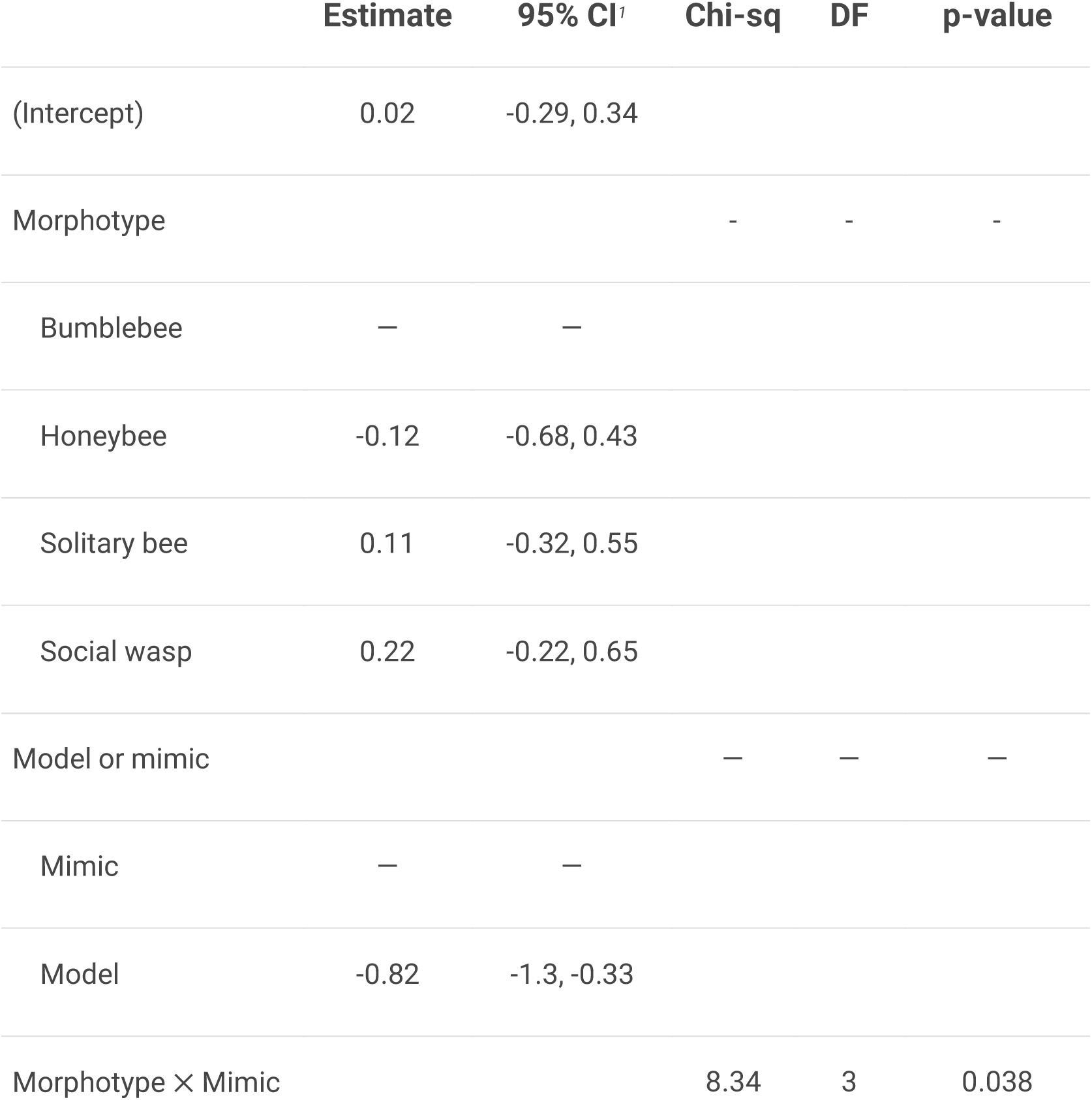
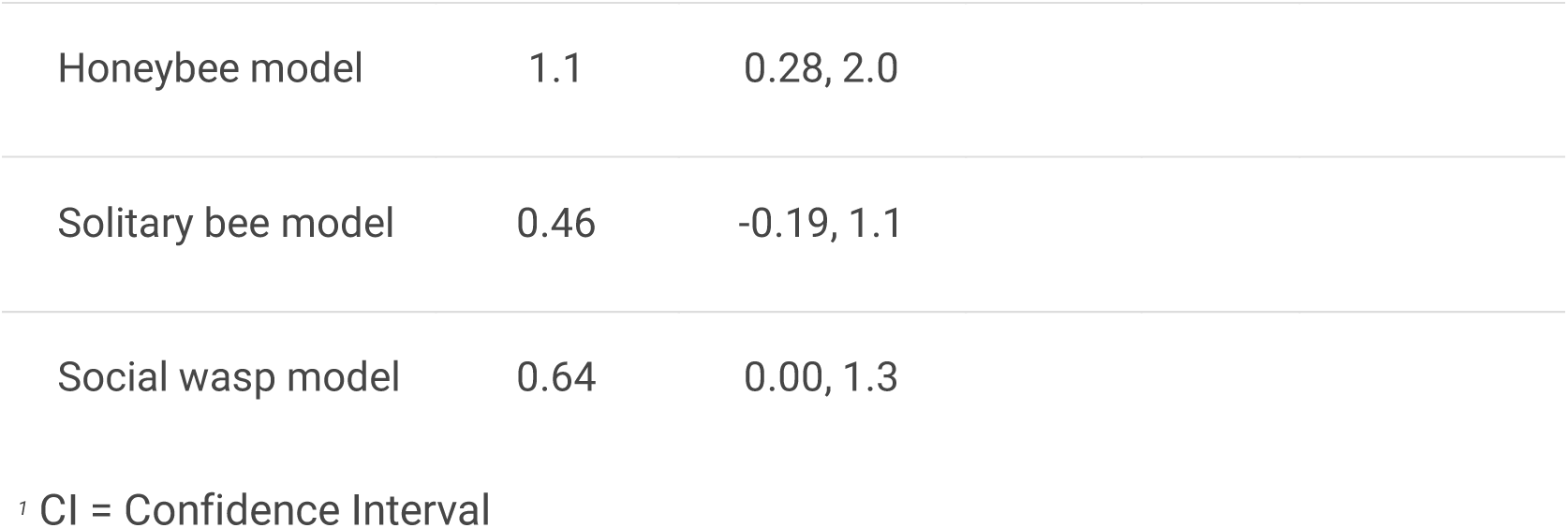
Linear mixed-effect model of flower yellowness against morphotype and “model or mimic”, with model-mimic pair as random effect.

|  | **Estimate** | **95% CI<sup>1</sup>** | **Chi-sq** | **DF** | **p-value** |
| --- | --- | --- | --- | --- | --- |
| (Intercept) | 0.02 | -0.29, 0.34 |  |  |  |
| Morphotype |  |  | - | - | - |
| Bumblebee | — | — |  |  |  |
| Honeybee | -0.12 | -0.68, 0.43 |  |  |  |
| Solitary bee | 0.11 | -0.32, 0.55 |  |  |  |
| Social wasp | 0.22 | -0.22, 0.65 |  |  |  |
| Model or mimic |  |  | — | — | — |
| Mimic | — | — |  |  |  |
| Model | -0.82 | -1.3, -0.33 |  |  |  |
| Morphotype × Mimic |  |  | 8.34 | 3 | 0.038 |

|  |  |  |
| --- | --- | --- |
| Honeybee model | 1.1 | 0.28, 2.0 |
| Solitary bee model | 0.46 | -0.19, 1.1 |
| Social wasp model | 0.64 | 0.00, 1.3 |

<sup>†</sup> CI = Confidence Interval

We then focused only on syrphids so that we could use phylogenetic information to ask whether mimics and non-mimics had diverged in preference, and whether body size as well as morphotype could explain flower colour preference.

Phylogenetic models of yellowness preference were practically indistinguishable from each other according to their DIC, including the model with no explanatory factors at all (delta DIC < 2.5). Across the phylogeny, syrphids of all morphotypes, mimics and non-mimics, preferred similar flower yellowness (see Table 3). In the model with technically the lowest DIC (delta ∼ 0.5), larger syrphids had a very slight tendency to prefer less yellow flowers which we regard as negligible.

**Table 3.** Table of fitted phylogenetic models of yellowness preference among syrphids, showing coefficients and ranked DIC values.

| (Intercept) | Morphotype | Head width | Morphotype<br>× head width | df | logLik | DIC | delta |
| --- | --- | --- | --- | --- | --- | --- | --- |
| 0.55 |  | + |  | 4 | -795.83 | 1607.17 | 0.00 |
| 2.37 | + | + | + | 12 | -793.67 | 1607.86 | 0.70 |
| 0.16 |  |  |  | 3 | -796.72 | 1608.04 | 0.88 |
| 0.52 | + | + |  | 8 | -795.41 | 1608.87 | 1.71 |
| 0.02 | + |  |  | 7 | -796.42 | 1609.60 | 2.44 |

In contrast, there were significant differences among groups in blueness preference (Linear Mixed model, X2=30.7, df=8, p<0.001). Solitary bee morphotypes and non-mimic hoverflies preferred flowers with the least blue. Honeybee and social wasp morphotypes were intermediate. Both bumblebee models and mimics preferred flowers with more blue than others.

When we excluded non-mimics and looked specifically at effects of mimicry and morphotype, there was no interaction between “model/mimic” and “morphotype”, and also no effect of “model/mimic”. This indicated that models tended to choose flowers with similar B values to those chosen by mimics. In contrast, there was a strong effect of morphotype (X2=26.0, df=3, p<0.0001), showing that different hymenopteran groups (and their syrphid mimics which follow their preferences) tend to choose flowers with distinctive blueness values.

The three phylogenetic models of blueness preference containing “morphotype” were indistinguishable from each other by their DIC (“morphotype only”, “morphotype + body size”, “morphotype X body size”; delta < 0.57). However, all were better than models without “morphotype” by > 4 DIC points (Table 6). This indicated that (as with the non-phylogenetic models above) morphotype, but not body size, was associated with blueness preference. Bumblebee mimics and, marginally, honeybee mimics had different blueness preferences from those of non-mimics, and from those of mimics of solitary bees and social wasps (see table 7).

**Table 4.**
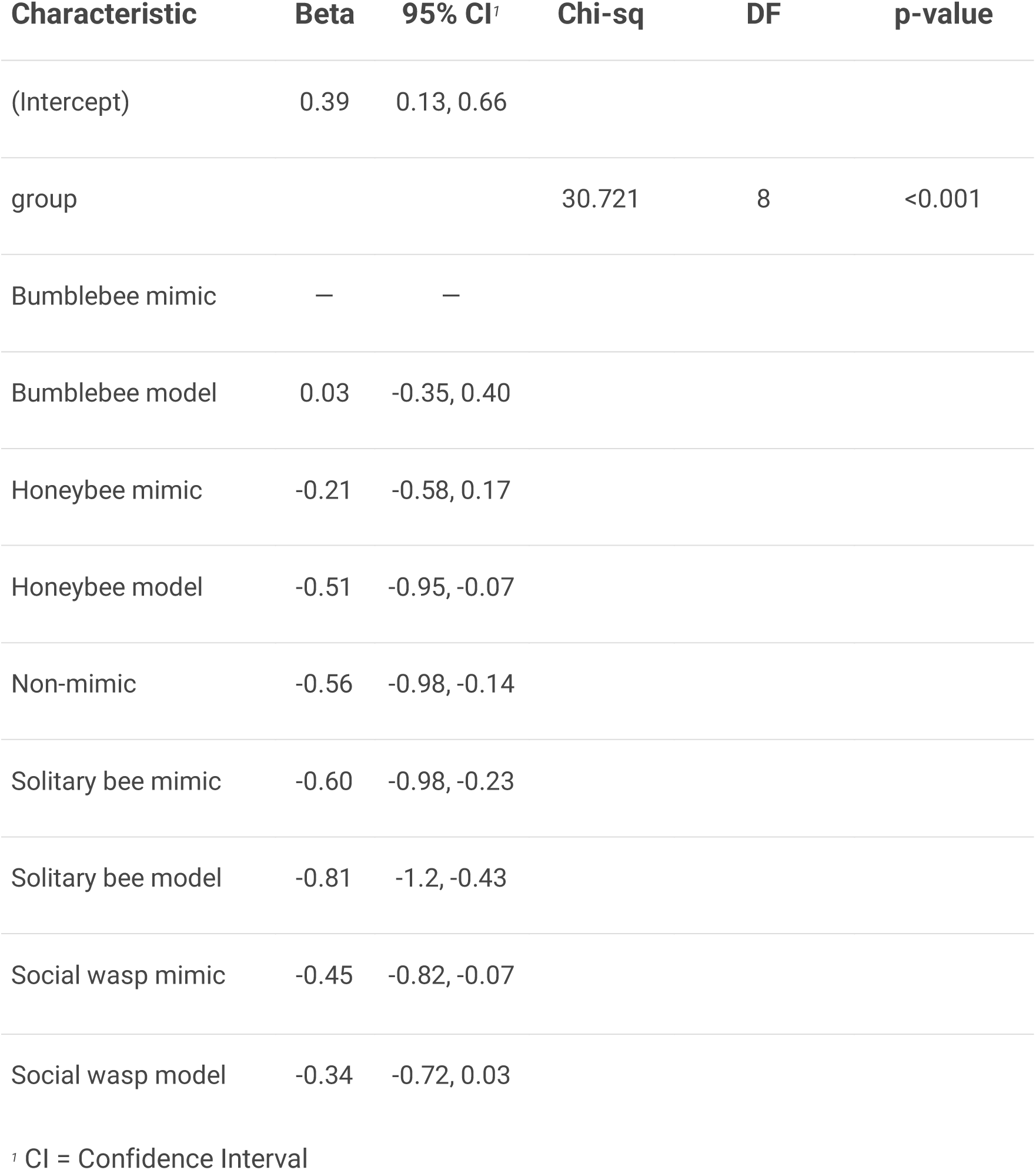
Linear mixed-effect model of flower blueness against morphotype, with species as random effect.

**Table 5.** Linear mixed-effect model of flower blueness against morphotype and “model or mimic”, with model-mimic pair as random effect.

|  | **Estimate** | **95% CI<sup>†</sup>** | **Chisq** | **DF** | **p-value** |
| --- | --- | --- | --- | --- | --- |
| Intercept) | 0.42 | 0.19, 0.66 |  |  |  |
| Morphotype |  |  | 21.31 | 3 | <0.0001 |
| Bumblebee | — | — |  |  |  |
| Honeybee | -0.34 | -0.72, 0.03 |  |  |  |
| Solitary bee | -0.69 | -0.98, -0.40 |  |  |  |
| Social wasp | -0.36 | -0.65, -0.08 |  |  |  |
| Model/mimic |  |  | 0.66 | 1 | 0.42 |
| Mimic | — | — |  |  |  |
| Model | -0.09 | -0.30, 0.13 |  |  |  |

<sup>†</sup> CI = Confidence Interval

**Table 6.**
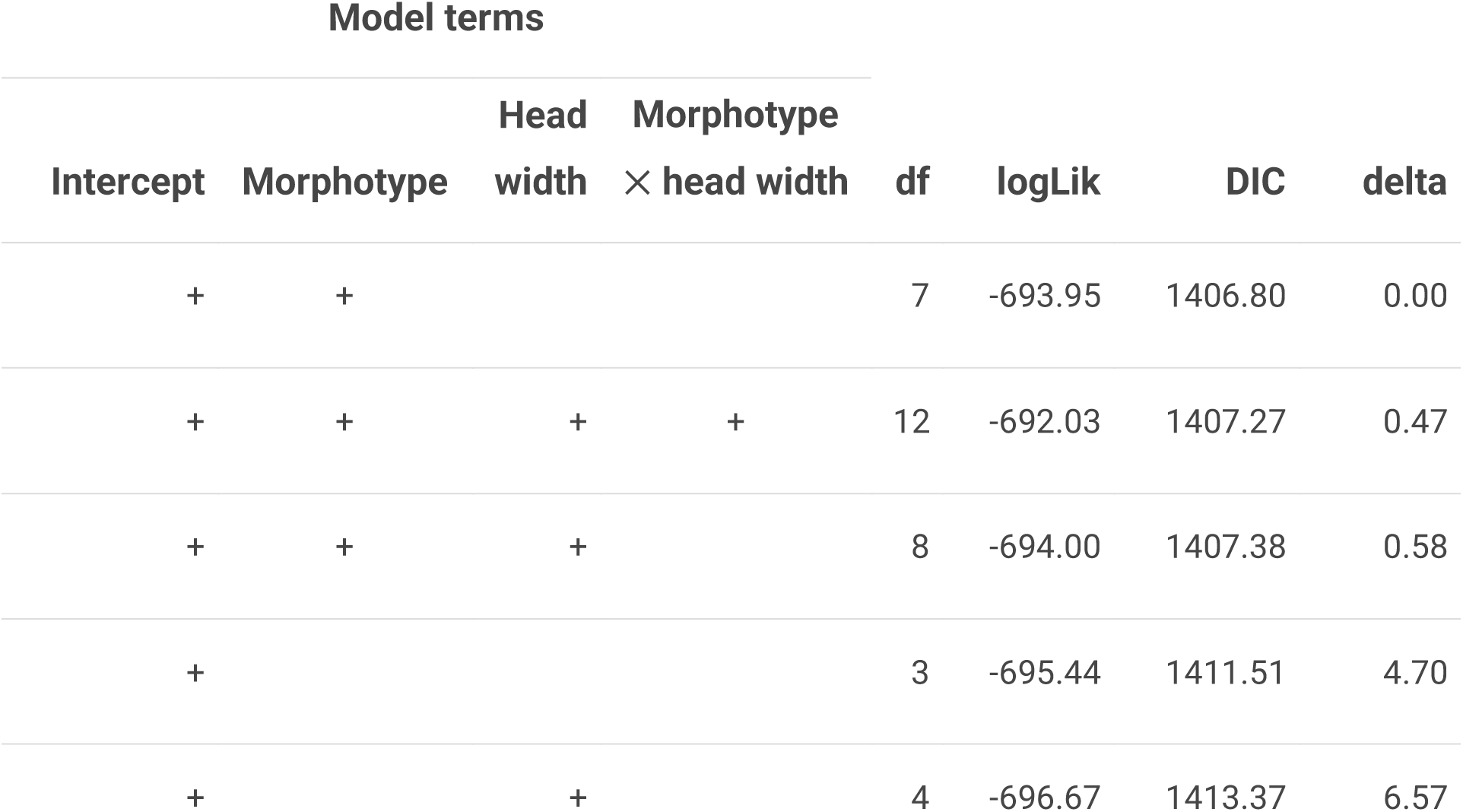
Table of fitted phylogenetic models of blueness preference among syrphids, showing coefficients and ranked DIC values.

**Table 7.**
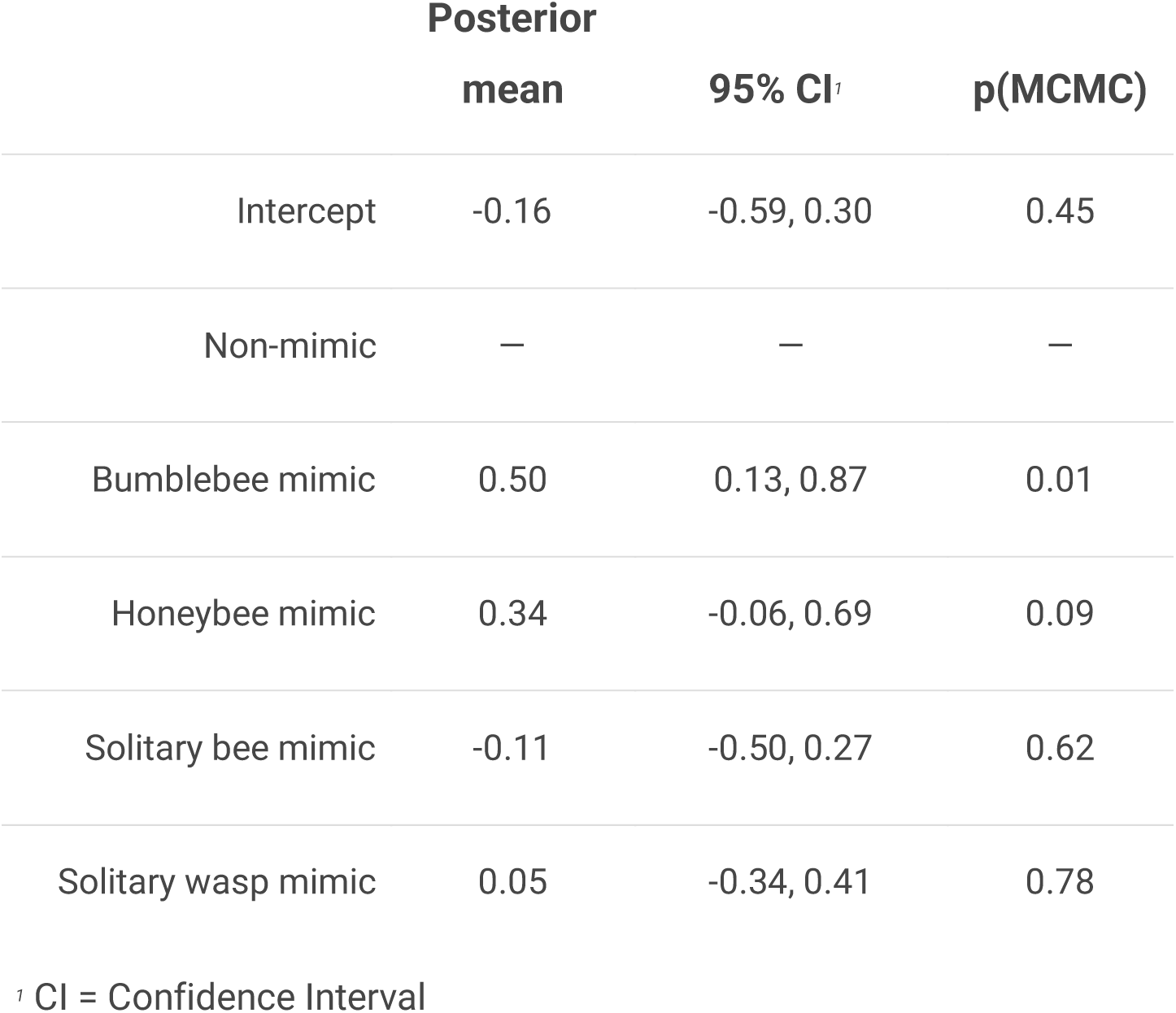
Table of fixed coefficients for top phylogenetic model of blueness preference among syrphids.

|  | Posterior<br>mean | 95% CI<sup>1</sup> | p(MCMC) |
| --- | --- | --- | --- |
| Intercept | -0.16 | -0.59, 0.30 | 0.45 |
| Non-mimic | — | — | — |
| Bumblebee mimic | 0.50 | 0.13, 0.87 | 0.01 |
| Honeybee mimic | 0.34 | -0.06, 0.69 | 0.09 |
| Solitary bee mimic | -0.11 | -0.50, 0.27 | 0.62 |
| Solitary wasp mimic | 0.05 | -0.34, 0.41 | 0.78 |

<sup>1</sup> CI = Confidence Interval

## Discussion

The “blueness” of flowers preferred by mimetic syrphids, and in particular bumblebee mimics, was statistically similar to that of their hymenopteran models - and in the case of bumblebee and honeybee mimics, was bluer than those preferred by non-mimetic syrphids. Bumblebee mimics such as *Volucella bombylans* have been documented as preferring blue flowers (Lunau, 2014), but to our knowledge this is the first systematic study to demonstrate a shift in flower colour preference in bumblebee mimics across the syrphid phylogeny. For their part, hymenopteran models showed differences in the blueness of flowers on which they tended to be photographed. Specifically, bumblebees, and to a lesser extent honeybees, were photographed on bluer flowers than other bees and wasps. This reflects a known preference towards blue and violet flowers as food sources (Giurfa et al., 1995; Gumbert, 2000; Lubbock, 1882; Raine et al., 2006) that probably indicates availability of floral nectar in suitably accessible locations for their generally long tongues (Raine & Chittka, 2007).

In yellowness preference, syrphid mimics also mostly tended to resemble their models, but their preferences were also similar to those of their non-mimic relatives. So, while this finding could be interpreted as behavioural mimicry of the models’ preferences by the syrphids, it could also simply reflect syrphid evolutionary ancestry, and/or dipteran ancestry generally. Dipterans, including hoverflies, have strong preferences towards yellow and white flowers in many species (Arnold et al., 2009) which again is probably related to accessibility of floral nectar for their relatively short tongues (see Lunau, 2014). Bumblebees were associated with less-yellow flowers than the other Hymenopterans, but bumblebee mimics did not follow this preference. Instead, they retained the preference of their non-mimic relatives and of all other syrphids for more-yellow flowers.

Why would syrphids mimicking social bees, in particular bumblebees, evolve away from their ancestral flower colour preferences to follow the flower blueness preferences of their models? It is difficult to escape the conclusion that their flower colour preferences have shifted to enhance their mimicry, either by providing a multicomponent stimulus (Rowe, 1999) or by bringing models and mimics into closer proximity to enhance protection (F. Gilbert, 2005). Bumblebee-mimicking syrphids include some of the most faithful of all syrphid mimics (F. Gilbert, 2005). Selection for high mimetic accuracy may be driven by both (1) large body size (Penney et al., 2012) and (2) mimicry of mildly noxious models such as bumblebees (Sherratt, 2002). On the one hand, Penney et al (2012) found that the largest syrphid mimics were the most faithful, and that small body size (and hence meagre reward for predators) was the best explanation for the occurrence of imperfect mimicry across species. Syrphids that show behavioural mimicry (mock stinging, wing-wagging etc.) tend to be those with large bodies and high-quality mimicry (although of wasps; Penney et al., 2014). While bumblebee mimics fit this description, in this study we found that body size did not affect preference for either flower blueness or yellowness in syrphids mimicking any kind of model.

On the other hand, Sherratt’s (2002) hypothesis holds that models such as wasps are so noxious to predators that any risk of misidentification confers protection even for poor quality mimics. For models such as bumblebees that are less dangerous to predators than wasps (F. Gilbert, 2005), the morphological “zone of protection” is more restricted, such that only near-perfect mimics are protected. Strong enough selection on mimetic accuracy may extend to selection for behavioural mimicry of flower colour preference. One interesting further question would be whether any such selection has been strong enough to distinguish finer-scale differences in colour preference among mimics of different bumblebee species. Fussell & Corbet (1992) report that bumblebee species “colour groups” (e.g. “brown”, “black with red abdominal band”) show distinct differences in flower usage, perhaps reflecting flowers with suitable corolla depths for their tongues (Brian, 1957; Faegri & van der Pijl, 1979). Flower colour loosely predicts corolla depth (Binkenstein et al., 2016) which may translate into colour preferences that mimics could exploit. Looking at the flower preferences (and tongue lengths) of the distinct morphs of polymorphic mimics such as *V. bombylans* and *Merodon equestris* might provide a good test of this hypothesis. Another natural further question is whether bumblebee-mimicking syrphids have evolved longer tongues to facilitate exploitation of “bee flowers” in the same way as bee-flies. Tongue length is particularly labile among hoverfly species independently of body size (F. S. Gilbert, 1985). *Rhingia*, a long-tongued genus of syrphids, although non-mimetic, are unusual amongst syrphids in preferring to visit red and blue flowers rather than yellow ones (Haslett, 1989; Robertson, 1928); more generally, flies that visit blue flowers often have elongate mouthparts (reviewed in Lunau, 2014).

Why mimic specifically the blueness and not the yellowness of bumblebee flower preferences? Some syrphids show an innate preference for yellow flowers that cannot be extinguished by training (Lunau et al., 2018), so it may be that the strong preference for yellow is stabilised by other evolutionary mechanisms and does not respond to selection for copying the preferences of models.

Ultimately, selection favouring mimicry arises from the visual systems of predators (Rowe, 1999; Speed & Turner, 1999). Internet photographs cannot possibly capture the range of colour involved in flower-pollinator-predator interactions (Stevens et al., 2007). Bees mostly have three kinds of photoreceptor: broadly, UV, blue and green (Briscoe & Chittka, 2001; Goldsmith, 1990), while syrphids and other flies have an effectively tetrachromatic system with an additional receptor in the UV range (An et al., 2018; Lunau, 2014). Birds also have potentially four receptors, but their additional receptor is in the red range (Hart & Hunt, 2007) which appears achromatic to insects - despite this, bird pollinators do not appear to show particular flower colour preferences that might exploit this private colour channel (Lunau et al., 2011). Whether avian predators pay more attention to the blueness of flowers visited by potential prey than to their yellowness remains a question to be investigated.

Of course, aspects of flowers other than colour are important in determining visitation by hoverflies, e.g. flower size (Sutherland et al., 1999) or height (Ssymank, 2001) and many others. Nevertheless, despite the obvious limitations of using internet photographs, we have managed to discern broad patterns consistent with behavioural mimicry of the flower-colour choices of hymenopteran models by syrphid mimics across the phylogeny, specifically in flower “blueness”. These patterns were most evident in bumblebee mimics, but were sufficiently strong that models and mimics generally were not statistically separable in their blueness preferences.

How diverse and multimodal traits interact to determine the effectiveness of antipredator defence is largely unknown (Kikuchi et al., 2023) - some traits combine with additive effects, others synergistic, and others antagonistic. Mimicry often involves multiple modalities (Rowe, 1999), such as morphology, colouration, behaviour, phenology and habitat use. Habitat-choice mimicry, such as we identify here, likely contributes to a “portfolio” of mimicry-enhancing traits in any given mimic species. Its costs and benefits likely depend upon factors such as predator-, model- and habitat-specific selection pressures (see e.g. Valkonen et al., 2012), competing allocation demands within the organism (Kikuchi et al., 2023), and how strong selection there is for mimetic accuracy in the first place (F. Gilbert, 2005; Penney et al., 2012). The hoverfly clade contains large numbers of origins of mimicry (Leavey et al., 2021) with members resembling diverse different models and employing multiple different modalities (Howarth & Edmunds, 2000; C. D. Moore & Hassall, 2016; Penney et al., 2014). We suggest that hoverflies are potentially an exceptional system in which to explore the evolution of multimodal mimicry, including the habitat-choice mimicry we have identified here.

## Acknowledgements

We thank M. Stevens for helpful advice on image processing; G. Ruxton for helpful comments on the manuscript; J. Boynton, K. Lester, E. Howard and other undergraduate students involved in this and similar projects over the years.

## Ethical Statement

All modification and usage of internet images in this study was for non-commercial research purposes (Levering, 1999, Jaszi, 2013). The research received ethical approval (in the form of a waiver) from the University of Hull Faculty Ethics Committee.

## Author contributions

JDJG conceived and co-designed the study, co-supervised data collection, conducted all data analysis and wrote the manuscript. JCJ collected all data and wrote an initial draft of the project as an undergraduate dissertation. LM co-designed the study, co-supervised data collection and edited the manuscript. FG provided data on models for different mimics, morphological data and phylogenetic information, and edited the manuscript.

## References

An, L., Neimann, A., Eberling, E., Algora, H., Brings, S., & Lunau, K. (2018). The yellow specialist: dronefly Eristalis tenax prefers different yellow colours for landing and proboscis extension. The Journal of Experimental Biology, 221(Pt 22), jeb184788.

Arnold, S. E. J., Savolainen, V., & Chittka, L. (2009). Flower colours along an alpine altitude gradient, seen through the eyes of fly and bee pollinators. Arthropod-Plant Interactions, 3(1), 27–43.

Atsumi, K., & Koizumi, I. (2017). Web image search revealed large-scale variations in breeding season and nuptial coloration in a mutually ornamented fish, *Tribolodon hakonensis*. Ecological Research, 32(4), 567–578.

Austen, E. J., Lin, S.-Y., & Forrest, J. R. K. (2018). On the ecological significance of pollen color: a case study in American trout lily (Erythronium americanum). Ecology, 99(4), 926– 937.

Bates, D. (2010). lme4 : linear mixed-effects models using S4 classes. http://cran.r-project.org/package=lme4. https://cir.nii.ac.jp/crid/1571698600750746240

Binkenstein, J., Stang, M., Renoult, J. P., & Schaefer, H. M. (2016). Weak correlation of flower color and nectar-tube depth in temperate grasslands. Journal of Plant Ecology, 10(2), rtw029.

Bot, S., & Van de Meutter, F. (2023). Hoverflies of Britain and North-west Europe: A photographic guide. Bloomsbury Wildlife.

Brian, A. D. (1957). Differences in the flowers visited by four species of bumble-bees and their causes. The Journal of Animal Ecology, 26(1), 71.

Briscoe, A. D., & Chittka, L. (2001). The evolution of color vision in insects. Annual Review of Entomology, 46(1), 471–510.

Ceccarelli, F. S. (2008). Behavioral mimicry in Myrmarachne species (Araneae, Salticidae) from North Queensland, Australia. The Journal of Arachnology, 36(2), 344–351.

de Buck, N. (1990). Bloembezoek en bestuivingsecologie van zweefvliegen (Diptera, Syrphidae) in het bijzonder voor België. Royal Belgian Institute of Natural Sciences.

Dlusski, G. M. (1984). Are dipteran insects protected by their similarity to stinging Hymenoptera. Byulleten’Moskovskogo Obshchestva Ispytatelei Prirody, Otdel Biologicheskii, 89, 25–40.

Faegri, K., & van der Pijl, L. (1979). The principles of pollination ecology - 3rd edition. https://agris.fao.org/search/en/providers/122621/records/647396dae0110688009805d2

Fussell, M., & Corbet, S. A. (1992). Flower usage by bumble-bees: A basis for forage plant management. The Journal of Applied Ecology, 29(2), 451.

Gilbert, F. (2005). The evolution of imperfect mimicry. Symposium-Royal Entomological Society of London. https://www.cabidigitallibrary.org/doi/pdf/10.1079/9780851998121.0000#page=244

Gilbert, F. S. (1985). Morphometric patterns in hoverflies (Diptera, Syrphidae). Proceedings of the Royal Society of London, 224(1234), 79–90.

Giurfa, M., Núñez, J., Chittka, L., & Menzel, R. (1995). Colour preferences of flower-naive honeybees. Journal of Comparative Physiology. A, Neuroethology, Sensory, Neural, and Behavioral Physiology, 177(3), 247–259.

Golding, Y. C., & Edmunds, M. (2000). Behavioural mimicry of honeybees (Apis mellifera) by droneflies (Diptera: Syrphidae: Eristalis spp.). Proceedings of the Royal Society of London. Series B: Biological Sciences, 267(1446), 903–909.

Golding, Y. C., & Ennos, A. R. (2005). Biomechanics and behavioural mimicry in insects. In A. Herrel, T. Speck, & N. P. Rowe (Eds.), Ecology and biomechanics: a mechanical approach to the ecology of animals and plants. Taylor & Francis.

Goldsmith, T. H. (1990). Optimization, constraint, and history in the evolution of eyes. The Quarterly Review of Biology, 65(3), 281–322.

Gumbert, A. (2000). Color choices by bumble bees (Bombus terrestris): innate preferences and generalization after learning. Behavioral Ecology and Sociobiology, 48(1), 36–43.

Hannah, L., Dyer, A. G., Garcia, J. E., Dorin, A., & Burd, M. (2019). Psychophysics of the hoverfly: categorical or continuous color discrimination? Current Zoology, 65(4), 483–492.

Hart, N. S., & Hunt, D. M. (2007). Avian visual pigments: characteristics, spectral tuning, and evolution. The American Naturalist, 169 Suppl 1(S1), S7–S26.

Haslett, J. R. (1989). Interpreting patterns of resource utilization: randomness and selectivity in pollen feeding by adult hoverflies. Oecologia, 78(4), 433–442.

Howarth, B., & Edmunds, M. (2000). The phenology of Syrphidae (Diptera): are they Batesian mimics of Hymenoptera? Biological Journal of the Linnean Society. Linnean Society of London, 71(3), 437–457.

Howarth, B., Edmunds, M., & Gilbert, F. (2004). Does the abundance of hoverfly (Syrphidae) mimics depend on the numbers of their hymenopteran models? Evolution; International Journal of Organic Evolution, 58(2), 367–375.

Ilse, D. (1949). Colour discrimination in the dronefly, Eristalis tenax. Nature, 163(4137), 255.

Kastinger, C., & Weber, A. (2001). Bee-flies (Bombylius spp., Bombyliidae, Diptera) and the pollination of flowers. Flora, 196(1), 3–25.

Kikuchi, D. W., Allen, W. L., Arbuckle, K., Aubier, T. G., Briolat, E. S., Burdfield-Steel, E. R., Cheney, K. L., Daňková, K., Elias, M., Hämäläinen, L., Herberstein, M. E., Hossie, T. J., Joron, M., Kunte, K., Leavell, B. C., Lindstedt, C., Lorioux-Chevalier, U., McClure, M., McLellan, C. F., … Exnerová, A. (2023). The evolution and ecology of multiple antipredator defences. Journal of Evolutionary Biology, 36(7), 975–991.

Kitamura, T., & Imafuku, M. (2015). Behavioural mimicry in flight path of Batesian intraspecific polymorphic butterfly Papilio polytes. Proceedings. Biological Sciences / The Royal Society, 282(1809), 20150483.

Kugler, H. (1950). Flower visitation of Eristalomyia tenax. Zeitschrift Für Vergleichende Physiologie, 3(4), 328–347.

Laitly, A., Callaghan, C. T., Delhey, K., & Cornwell, W. K. (2021). Is color data from citizen science photographs reliable for biodiversity research? Ecology and Evolution, 11(9), 4071–4083.

Leavey, A., Taylor, C. H., Symonds, M. R. E., Gilbert, F., & Reader, T. (2021). Mapping the evolution of accurate Batesian mimicry of social wasps in hoverflies. Evolution; International Journal of Organic Evolution, 75(11), 2802–2815.

Leighton, G. R. M., Hugo, P. S., Roulin, A., & Amar, A. (2016). Just Google it: assessing the use of Google Images to describe geographical variation in visible traits of organisms. Methods in Ecology and Evolution / British Ecological Society, 7(9), 1060–1070.

Lenth, R. V. (2023). emmeans: Estimated Marginal Means, aka Least-Squares Means. https://CRAN.R-project.org/package=emmeans

Lubbock, J. (1882). Ants. Bees, and wasps. Kegan Paul, Trench & Company.

Lunau, K. (2014). Visual ecology of flies with particular reference to colour vision and colour preferences. Journal of Comparative Physiology. A, Neuroethology, Sensory, Neural, and Behavioral Physiology, 200(6), 497–512.

Lunau, K., An, L., Donda, M., Hohmann, M., Sermon, L., & Stegmanns, V. (2018). Limitations of learning in the proboscis reflex of the flower visiting syrphid fly Eristalis tenax. PloS One, 13(3), e0194167.

Lunau, K., Papiorek, S., Eltz, T., & Sazima, M. (2011). Avoidance of achromatic colours by bees provides a private niche for hummingbirds. The Journal of Experimental Biology, 214(Pt 9), 1607–1612.

Mappes, J., & Alatalo, R. V. (1997). Batesian Mimicry and Signal Accuracy. Evolution; International Journal of Organic Evolution, 51(6), 2050.

Moore, C. D., & Hassall, C. (2016). A bee or not a bee: an experimental test of acoustic mimicry by hoverflies. Behavioral Ecology: Official Journal of the International Society for Behavioral Ecology, 27(6), 1767–1774.

Moore, M. P., Lis, C., Gherghel, I., & Martin, R. A. (2019). Temperature shapes the costs, benefits and geographic diversification of sexual coloration in a dragonfly. Ecology Letters, 22(3), 437–446.

Morse, D. H. (1979). Prey capture by the crab spider Misumena calycina (Araneae: Thomisidae). Oecologia, 39(3), 309–319.

Penney, H. D., Hassall, C., Skevington, J. H., Abbott, K. R., & Sherratt, T. N. (2012). A comparative analysis of the evolution of imperfect mimicry. Nature, 483(7390), 461–464.

Penney, H. D., Hassall, C., Skevington, J. H., Lamborn, B., & Sherratt, T. N. (2014). The relationship between morphological and behavioral mimicry in hover flies (Diptera: Syrphidae). The American Naturalist, 183(2), 281–289.

Raine, N. E., & Chittka, L. (2007). The adaptive significance of sensory bias in a foraging context: floral colour preferences in the bumblebee Bombus terrestris. PloS One, 2(6), e556.

Raine, N. E., Ings, T. C., Dornhaus, A., Saleh, N., & Chittka, L. (2006). Adaptation, genetic drift, pleiotropy, and history in the evolution of bee foraging behavior. In Advances in the Study of Behavior (Vol. 36, pp. 305–354). Elsevier.

R Core Team. (2023). R: A Language and Environment for Statistical Computing. R Foundation for Statistical Computing. https://www.R-project.org/

Rettenmeyer, C. W. (1970). Insect mimicry. Annual Review of Entomology, 15(1), 43–74.

Robertson, C. (1928). Flowers and insects: XXV. Ecology, 9(4), 505–526.

Rodríguez-Gasol, N., Avilla, J., Alegre, S., & Alins, G. (2019). Sphaerophoria rueppelli adults change their foraging behavior after mating but maintain the same preferences to flower traits. BioControl (Dordrecht, Netherlands), 64(2), 149–158.

Rowe, C. (1999). Receiver psychology and the evolution of multicomponent signals. Animal Behaviour, 58(5), 921–931.

Ruxton, G. D. (2009). Non-visual crypsis: a review of the empirical evidence for camouflage to senses other than vision. Philosophical Transactions of the Royal Society of London. Series B, Biological Sciences, 364(1516), 549–557.

Sherratt, T. N. (2002). The evolution of imperfect mimicry. Behavioral Ecology: Official Journal of the International Society for Behavioral Ecology, 13(6), 821–826.

Skevington, J. H., Locke, M. M., Young, A. D., Moran, K., Crins, W. J., & Marshall, S. A. (2019). Field guide to the flower flies of northeastern north America. Princeton University Press.

Smith, J., & Joost, R. (2012). Digital Imaging Projects. In GIMP for Absolute Beginners (pp. 131–166). Apress.

Speed, M. P., & Turner, J. R. G. (1999). Learning and memory in mimicry: II. Do we understand the mimicry spectrum? Biological Journal of the Linnean Society. Linnean Society of London, 67(3), 281–312.

Ssymank, A. (1989). Das Vegetationsmosaik eines Waldgebietes der Schwarzwaldvorbergzone und seine funktionale Bedeutung für blütenbesuchende Insekten-unter besonderer Berücksichtigung der Syrphidae (Diptera): Text: mit 12 pflanzensoziologischen Tab., 22 phänologischen Tab., 7 biozönologischen und blütenbiologischen Tab. und weiteren 33 Tab. im Text.

Ssymank, A. (2001). Vegetation and flower-visiting insects in cultivated landscapes. Bonn Bad Godesb, 64, 513.

Stevens, M., Párraga, C. A., Cuthill, I. C., Partridge, J. C., & Troscianko, T. S. (2007). Using digital photography to study animal coloration. Biological Journal of the Linnean Society. Linnean Society of London, 90(2), 211–237.

Sutherland, J. P., Sullivan, M. S., & Poppy, G. M. (1999). The influence of floral character on the foraging behaviour of the hoverfly, Episyrphus balteatus. Entomologia Experimentalis et Applicata, 93(2), 157–164.

Szucsich, N. U., & Krenn, H. W. (2000). Morphology and function of the proboscis in Bombyliidae (Diptera, Brachycera) and implications for proboscis evolution in Brachycera. Zoomorphology, 120(2), 79–90.

Thompson, F. (1991). The flower fly genus Ornidia (Diptera: Syrphidae). Proceedings of the Entomological Society of Washington, 93(2), 248–260.

Tkalcic, M., & Tasic, J. F. (2003). Colour spaces: perceptual, historical and applicational background. The IEEE Region 8 EUROCON 2003. Computer as a Tool, Ljubljana, Slovenia. https://ieeexplore.ieee.org/abstract/document/1248032

Valkonen, J. K., Nokelainen, O., Niskanen, M., Kilpimaa, J., Björklund, M., & Mappes, J. (2012). Variation in predator species abundance can cause variable selection pressure on warning signaling prey. Ecology and Evolution, 2(8), 1971–1976.

Dlusskii GM (1984) [Are Diptera protected by their similarity to stinging Hymenoptera?] Byulleten Moskovskogo Obshchestva Ispytatelei Prirody Otdel Biologicheskii 89(5): 25–40 (in Russian)

Gilbert FS (1985) Morphometric patterns in hoverflies (Diptera, Syrphidae). Proceedings of the Royal Society of London B Biological Sciences 224: 79–90

Szucsich NU & Krenn HW (2000) Morphology and function of the proboscis in Bombyliidae (Diptera, Brachycera) and implications for proboscis evolution in Brachycera. Zoomorphology 120: 79–90

