## Supplementary material for "Internet images reveal bumblebee-mimicking hoverflies have evolved to follow flower colour preferences of their models": 2024 Syrphid behavioural mimicry - results and code.Rmd

````
---
title: "Internet images reveal bumblebee-mimicking hoverflies have evolved to follow flower colour preferences of their models - Results and code"
output:
  html_document:
    df_print: paged
---

```{r preliminaries, include=F , echo=F}

##  - RStudio's default CRAN mirror doesn't appear to have some packages
r <- getOption("repos")
    r["CRAN"] <- "https://cloud.r-project.org"
    options(repos = r)

### Check for missing packages and install them if necessary
    # https://stackoverflow.com/questions/4090169/elegant-way-to-check-for-missing-packages-and-install-them
    # - installs the package if you don't already have it
using<-function(...) {
    libs<-unlist(list(...))
    req<-unlist(lapply(libs,require,character.only=TRUE)) # install the ones you have
    need<-libs[req==FALSE] # list of ones you didn't have
    if(length(need)>0){ 
        install.packages(need) # install anything you didn't previously have
        lapply(need,require,character.only=TRUE) # load them up after installing
    }
}

# run this for the packages we need
using("foreach", "lme4", "nlme", "MCMCglmm", "phytools", "Hmisc", "vegan", "metafor", "ggplot2", "patchwork", "tidyr", "tidyverse", "magrittr", "emmeans","MuMIn","ggtree", "tidytree", "viridis", "gtsummary", "gt")


```

```{r functions, include = F, echo = F}

# function to get the model species from the dataset for a given mimic
get_model_species <- function(x) {
    unique(dat$species[grep(unique(dat$pair2[dat$species==x][1]), tolower(dat$species))])
}

# function to extract the subset of data for the given model species set
bind_model_species <- function(x) {

dat.mimicx <- filter(dat, species == x)
dat.modelx <- filter(dat, species %in% get_model_species(x))

dat.x <- 
  rbind.data.frame(dat.mimicx, dat.modelx) %>% 
    mutate(mimsp = rep(x, nrow(dat.mimicx)+nrow(dat.modelx)))

dat.x

}

# function to create a model-mimic group
group_model_species <- function(x) do.call(rbind.data.frame, lapply(x, bind_model_species))

# function to scale any variable between 0 and 1
fractionize <- function(x) sapply(x, function(y) ((y-min(x, na.rm=T)))/(max(x, na.rm=T)-min(x, na.rm=T)))

## function to build a star phylogeny from a list of species
buildtree <- function(X) {

test.tree <- read.tree(text=paste('(', 
					paste(X, collapse=','),
					');', sep=''))
test.tree
}


```


```{r data, include=F , echo=F}

nrow(dat <- read.csv('Table S3 main data.csv'))
dat$id <- dat$species

dat$species <- trimws(gsub('[0-9]','', dat$id)) # remove any trailing spaces and numbers

dat$species[grep('bombylans', dat$species)] <- 'Volucella_bombylans' #trim variant info


# morphotype and mimic stored in raw data as one single column "group" -
# transform "group" column into "morphotype" column and "mimic" column
dat$mtype <- factor(sub('mod|mim','', dat$group), levels=c('bbee','hbee','sbee','swasp','non'))

## column denoting mimic or model
dat$mimic <- sapply(as.character(dat$group), function(x) substr(x, nchar(x)-2, nchar(x)))


dat$species <- capitalize(gsub(' ','_', dat$species)) # ensure species name format consistent
dat$genus <- capitalize(sub('_.*','',dat$species)) # Create genus column
dat$genus[dat$genus=='Pyrophaena'] <- 'Platycheirus' ## synonym
dat$genus[dat$genus=='Anasiymia'] <- 'Anasimyia'

dat$pair2<-tolower(sub(" ","_", trimws(dat$pair2))) # ensure model string format consistent

```


```{r add morphological data, include=F, echo=F}
### import mean body size (thorax width) data for species

morph <- read.csv('Table S2 head width.csv') %>% 
            transmute(binomial=species,
                   meanhw=`head.width..mm.`) 

```


```{r PCA on log-transformed data, include=F , echo=F}

## data are strongly left skewed, so transform with log(constant - x)
dat.trans <- transmute(dat,
                      tr = -log(255-r),
                      tg = -log(255-g),
                      tb = -log(255-b))
    ## multiply by -1 as negatively related to original value

## Fit PCA
pca1 <- prcomp(dat.trans, scale=T)

## Add PCA axes to dataset
dat$pca1 <- pca1$x[,1] 
            ## since pca1 is correlated 0.70 with both R and G
dat$pca2 <- -pca1$x[,2]
            ## since pca2 is correlated -0.97 with B
            ## need pca2 to be positive "blueness", hence multiply by -1

```

```{r import phylogeny, include=F , echo=F}

tree <- read.nexus('Supplementary file S4 - phylogeny.nex') 
# Ntip(tree) # 251 tips [for troubleshooting]

```

```{r sync phylogeny for model-mimic differences, include=F , echo=F}

 ## dataset of mimics only, no models
dat.mimics <- subset(dat, mimic=='mim')
 # length(intersect(dat.mimics$genus, tree$tip.label)) # 25 [for troubleshooting]
length(unique(dat.mimics$genus))
 # setdiff(dat.mimics$genus, tree$tip.label) # 0
 
dat.mimics$group <- factor(dat.mimics$group)

# restrict genus tree
tree.dat <- drop.tip(tree, tip=which(!tree$tip.label %in% dat.mimics$genus))
 # Ntip(tree.dat) # 25 [for troubleshooting]

## add in species trees as star phylogenies
gentable <- table(subset(dat.mimics, !duplicated(species))$genus)
genx <- names(gentable)[which(gentable>1)]
sppx <- capitalize(unique(dat.mimics$species[dat.mimics$genus%in%genx]))


## for functions used below see "functions" chunk above


# create a star phylogeny for each genus with >1 member
for (i in genx)  {
	 assign(i, buildtree(unique(dat.mimics$species[dat.mimics$genus==i])))
	 }
 
# add these to the genus-level phylogeny
for (i in genx) {
	 tree.dat <- bind.tree(tree.dat, get(i), where=grep(paste('^',i,'$',sep=''), tree.dat$tip.label))
		 }

geny <- tree.dat$tip.label[-grep('_', tree.dat$tip.label)]

for(i in geny) tree.dat$tip.label[tree.dat$tip.label==i] <- unique(dat.mimics$species[dat.mimics$genus==i])
  
## Remove node labels
tree.dat$node.label<-NULL

## Apply Grafen's transformation
tree.dat<-compute.brlen(tree.dat,power=0.8)

# Ntip(tree.dat) # 40 [for troubleshooting]

```


```{r Phylogeny to match dat.syrph - all syrphids inc non mimics, include=F, echo=F}

  
# dataset of all syrphids, incl non mimics
dat.syrph <- subset(dat, mimic %in% c('mim','non'))
dat.syrph$animal <- dat.syrph$species
dat.syrph$group <- factor(dat.syrph$group)

tree.syrph <- drop.tip(tree, tip=which(!tree$tip.label %in% dat.syrph$genus))
        #  29 tips
gentable <- table(subset(dat.syrph, !duplicated(species))$genus)
genx <- names(gentable)[which(gentable>1)]
sppx <- capitalize(unique(dat.syrph$species[dat.syrph$genus%in%genx]))


## for functions used below see "functions" chunk above
  
# create a star phylogeny for each genus with >1 member
for (i in genx)  {
	 assign(i, buildtree(unique(dat.syrph$species[dat.syrph$genus==i])))
	 }

# add these to the genus-level phylogeny
for (i in genx) {
	 tree.syrph <- bind.tree(tree.syrph, get(i), where=grep(paste('^',i,'$',sep=''), tree.syrph$tip.label))
		 }

geny <- tree.syrph$tip.label[-grep('_', tree.syrph$tip.label)]

for(i in geny) tree.syrph$tip.label[tree.syrph$tip.label==i] <- unique(dat.syrph$species[dat.syrph$genus==i])
  
# remove all node labels
tree.syrph$node.label<-NULL

## apply grafen's transformation to branch lengths which are arbitrary
tree.syrph<-compute.brlen(tree.syrph,power=0.8)
  
```

The 45 syrphids in the dataset, their mean colour preferences, and those of their models, can be seen in Figure 1.


```{r calculate median flower colour per species, include = F, echo = F}

## Calculate median flower colour for each syrphid species
mean_colour <-
dat.syrph %>% 
  group_by(group, species) %>% 
    summarize(median_r = median(r/255), median_g = median(g/255), median_b = median(b/255)) %>%       
                mutate(hex = rgb(median_r, median_g, median_b),
                groupcol = c('blue','brown','black','orange','yellow')[as.numeric(group)]) %>% 
             slice(order(factor(species, levels=tree.syrph$tip.label))) %>% 
        select(species, groupcol, hex) 

## For each syrphid, calculate median flower colour for models
mean_model_r <-  sapply(mean_colour$species, 
                       function(x) {
                         temp <- bind_model_species(x)
                          return(median(temp$r[temp$mimic=='mod']/255, na.rm=T))
                       })

mean_model_g <-  sapply(mean_colour$species, 
                       function(x) {
                         temp <- bind_model_species(x)
                          return(median(temp$g[temp$mimic=='mod']/255, na.rm=T))
                       })
  

mean_model_b <-  sapply(mean_colour$species, 
                       function(x) {
                         temp <- bind_model_species(x)
                          return(median(temp$b[temp$mimic=='mod']/255, na.rm=T))
                       })
  
mean_colour$modelr <- mean_model_r[match(mean_colour$species, names(mean_model_r))]
mean_colour$modelg <- mean_model_g[match(mean_colour$species, names(mean_model_g))]
mean_colour$modelb <- mean_model_b[match(mean_colour$species, names(mean_model_b))]

mean_colour$model_avg_rgb <- with(mean_colour, rgb(modelr, modelg, modelb))

## join this to head width data
mean_colour <-
  mean_colour %>% 
    left_join(morph, by = join_by(species == binomial), keep = T)


df <- mean_colour

#confirm no NA values
table(is.na(df$meanhw))
#FALSE 
#   45 

```


```{r add mean head width back in to original data, include=F, echo=F}

## get HW from df
dat.syrph$meanhw <- df$meanhw[match(dat.syrph$species, df$species)]


```


(Paste phylogeny figure here after manually adding labels and species images)

Figure 1. Syrphid phylogeny used in this study (with arbitrarily scaled branch lengths) showing morphotype, body size, and the flower colour preferences of mimics and models, as assessed by internet images depicting the species visiting a flower. Photos of representative species are given, but are not necessarily part of the dataset.


```{r Syrphid phylogeny showing colour prefs, include=F, echo=F, message=F, warnings=F}

## This chunk creates the base phylogeny figure but it is then exported to 
## MS Powerpoint and labels added manually, hence it is not included in output here

meanhw_frac <- 1-fractionize(df$meanhw)

## calculate viridis colour scale for body size
meanhw_col <- sapply(meanhw_frac, function(x) viridis(100)[round(x*99+1)])
        ## generates 100 viridis colours, chooses
        ## the nth colour based on the %ile head width
        ## calculated using fractionize() (see functions above)

df$meanhw_col <- meanhw_col

# Figure produced using tidytree
# code from https://stackoverflow.com/questions/78618050/unable-colour-the-label-tips-of-my-ggtree

x <- full_join(as_tibble(tree.syrph),
  df,
  by = c("label" = "species")
)
tree2 <- as.treedata(x)


p<- ggtree(tree2) + 
  geom_tiplab(size=2, offset=0.25) + 
    geom_tippoint(pch=15, size=3.5, position=position_nudge(x=0.00), aes(col=groupcol)) +
    geom_tippoint(pch=15, size=3.5, position=position_nudge(x=0.05), aes(col=meanhw_col)) +
  
    geom_tippoint(pch=15, size=5.5, position=position_nudge(x=0.15), aes(col=grey(0.5))) + 
    geom_tippoint(pch=15, size=3.5, position=position_nudge(x=0.15), aes(col=model_avg_rgb)) +
    geom_tippoint(pch=15, size=5.5, position=position_nudge(x=0.19), aes(col=grey(0.5))) +  
    geom_tippoint(pch=15, size=3.5, position=position_nudge(x=0.19), aes(col=hex)) +
    scale_colour_identity() +
    xlim(0, 2)
    
p

```


Two axes explained 83% of the variance in the colour of flowers photographed being visited by models or mimics.  Axis 1 (49% of the variance) was aligned strongly and identically with red (71%) and green (70%) and will be known henceforth as yellowness. Axis 2 (34% of the variance) was exclusively aligned with blue (98%) and we refer to this axis as blueness.


```{r, include=F , echo=F}

summary(pca1)
pca1

```


Scores along the yellowness and blueness axes for different morphotypes (models and mimics) are shown in Figure X.

Figure X. Blueness (PCA2) plotted against yellowness (PCA1) for models and mimics of different morphotypes, with means and 95% confidence ellipses shown in solid colours. Inset shows RGB colours for the whole dataset, with representative extremes marked for comparison.


```{r Figure X: PCA model-mimic biplots for different morphotypes, include=T, echo=F, message=F, warnings=F}

### Panels and inset are produced separately and exported to MS Powerpoint where they are
### combined manually.

medianpca1 <- dat %>% 
     mutate(mtype=recode(mtype, 
         bbee = 'Bumblebee',
         hbee = 'Honeybee',
         sbee = 'Solitary bee',
         swasp = 'Social wasp',
         non = 'Non-mimic'),
         mimic = recode(mimic, 
         mim = 'Mimics',
         mod = 'Models',
         non = 'Non-mimics')) %>% 
    group_by(mtype, mimic) %>% 
    summarise(median_val = median(pca1))

medianpca2 <- dat %>% 
     mutate(mtype=recode(mtype, 
         bbee = 'Bumblebee',
         hbee = 'Honeybee',
         sbee = 'Solitary bee',
         swasp = 'Social wasp',
         non = 'Non-mimic'),
         mimic = recode(mimic, 
         mim = 'Mimics',
         mod = 'Models',
         non = 'Non-mimics')) %>% 
    group_by(mtype, mimic) %>% 
    summarise(median_val = median(pca2))

fig2 <- 
dat %>% 
  mutate(mtype=recode(mtype, 
         bbee = 'Bumblebee',
         hbee = 'Honeybee',
         sbee = 'Solitary bee',
         swasp = 'Social wasp',
         non = 'Non-mimic'),
         mimic = recode(mimic, 
         mim = 'Mimics',
         mod = 'Models',
         non = 'Non-mimics')) %>% 
  ggplot(aes(y=pca2, x=pca1, col = mimic)) +
        geom_hline(aes(yintercept=0), linetype=2) +
        geom_vline(aes(xintercept=0), linetype=2) +
        ylab('Blueness (PCA2)') +
        xlab('Yellowness (PCA1)') +
      geom_point() +
        geom_hline(data=medianpca2, aes(yintercept=median_val, col=mimic)) +
        geom_vline(data=medianpca1, aes(xintercept=median_val, col=mimic)) +
        stat_ellipse() +
        facet_wrap(~mtype) +
        theme_classic() +
  theme(
   strip.background = element_rect(
     color=NA, fill=NA, size=0
     )
   )
 fig2

 ## we want to be able to generate the colour of points at the 
## extreme ends of the PCA distribution for comparison
 
## Here's a way of backcalculating the original input data from a PCA
## from https://stats.stackexchange.com/questions/229092/how-to-reverse-pca-and-reconstruct-original-variables-from-several-principal-com

## reconstructing original data
nComp <- 2
Xpca <- pca1
Xhat <- Xpca$x[,1:nComp] %*% t(Xpca$rotation[,1:nComp]) 
mu <- colMeans(dat.trans)
Xhat <- scale(Xhat, center = -mu, scale = FALSE)
X1 <- pmax(255-exp(-Xhat),0) ## backtransform to original RGB values (they had pvsly been logged)
                      ## coercing a few negatives to zero where necessary

## predict new RGB data from hypothetical PCA values based on the range of data

## Extremes of PCA
pca1_range <- range(dat$pca1)
pca2_range <- range(dat$pca2)
x <- c(pca1_range[1], pca1_range[2], 0, pca1_range[1], pca1_range[2])
y <- c(pca2_range[1], pca2_range[1], 0, pca2_range[2], pca2_range[2])

## Backtransform these to RGB
comp <- data.frame(x, y)
Xhat <- as.matrix(comp) %*% t(Xpca$rotation[,1:nComp]) # use "comp" instead of pca scores
mu <- colMeans(dat.trans)
Xhat <- scale(Xhat, center = -mu, scale = FALSE)
X <- pmax(255-exp(-Xhat), 0) # coerce negatives to zero

# Base plot
plot(-dat$pca2~dat$pca1,
        xlim=c(-4.5, 4.5), ylim=c(-2.5, 4.5), type='n',
              		xlab='yellowness (PCA1)',
    		ylab='blueness (PCA2)', 
)

# grey background
rect(par("usr")[1], par("usr")[3],
     par("usr")[2], par("usr")[4],
     col = "#dddddd") # Color

# points from original data (not original colour; reconstructed from PCA)
points(dat$pca2~dat$pca1,
        xlim=c(-4.5, 4.5), ylim=c(-2.5, 4.5),
    		    cex=1.5, pch=16, # col='grey',
            col = apply(X1, 1, function(x) 
                  do.call(rgb, 
                        list(x[1]/255, x[2]/255, x[3]/255)
                        )
                  )
)

## add comparison points at extremes of the PCA distribution
x1 <- c(-4, 4, 0, -4, 4)
y1 <- c(4, 4, 0, -2, -2) 
 points(x1, y1, pch=21, cex=3,
       bg = apply(X, 1, function(x) 
                  do.call(rgb, 
                        list(x[1]/255, x[2]/255, x[3]/255)
                        )
                  )
       )

## code for adding this schematic as an inset to the 
## main figure which is drawn using ggplot/facet_wrap
 
## For submission I have just pasted it in manually 
 ## upon acceptance will be added here and formatted consistently
 
#library(gtable)
#library(gridExtra)
#library(grid)
#
#pg <- ggplotGrob(fig2)
#qg <- ggplotGrob(fig2inset)
#
#pl <- gtable_filter(pg, 'panel', trim=F)$layout
#pg <- gtable_add_grob(pg, qg, t=max(pl$t), l=max(pl$l))
#
#grid.newpage()
#grid.draw(pg)

```


```{r data subset excluding nonmimics, include=F , echo=F}
## data subset excluding nonmimics
dat.mimics.models <- droplevels(subset(dat, mimic!='non'))

```


Hymenopteran models of different morphotypes differed in the yellowness of the flowers they chose to visit, while syrphids (mimics and non-mimics) all chose flowers of similar yellowness (Linear mixed model, X2=32.5, df=8, p<0.0001, Table 1, Figure 3). When we examined effects of mimicry and morphotype independently, excluding non-mimic hoverflies, there was a clear "mimic/model" x morphotype interaction, X2=8.45, df=3, p=0.037, Table 2).  Bumblebees chose less yellow flowers than other models, which did not differ from each other (honeybee, solitary bee and social wasp). In contrast, all mimic syrphids chose a similar yellowness, which was also similar to that chosen by honeybees, solitary bees and social wasps, but was higher than the yellowness chosen by bumblebees.


Table 1. Linear mixed-effect model of flower yellowness against morphotype, with species as random effect.

```{r Table 1, include=T, echo=T}
lmer1 <- lmer(pca1 ~ group + (1 | species),  data=dat) 
lmer2 <- lmer(pca1 ~ 1 + (1 | species),  data=dat) 

anova(lmer1, lmer2) # sig, obvs

emmeans(lmer1, "group")


# Table 1
lmer1 %>% tbl_regression(intercept=T)  # Manually add LRT values to this
```

Table 2. Linear mixed-effect model of flower yellowness against morphotype and "model or mimic", with model-mimic pair as random effect.

```{r Table 2 lmer analysis - PCA1 (yellow) ~ "model group" * "model/mimic" removing nonmimics, include=T , echo=F}
#Does "who you look like" * "model or mimic" affect flower colour preference?
#PCA1

lmer1 <- lmer(pca1 ~ mtype * mimic + (1 | pair_species),  data=dat.mimics.models) 
lmer2 <- lmer(pca1 ~ mtype + mimic + (1 | pair_species), data=dat.mimics.models)
lmer3 <- lmer(pca1 ~          mimic + (1 | pair_species), data=dat.mimics.models)
lmer4 <- lmer(pca1 ~ mtype         + (1 | pair_species), data=dat.mimics.models)
lmer5 <- lmer(pca1 ~   1            +  (1 | pair_species), data=dat.mimics.models)

anova(lmer1, lmer2) ## Interaction significant p=0.037
anova(lmer2, lmer3) ## Effect of morphotype
anova(lmer2, lmer4) ## Effect of "model or mimic"
anova(lmer4, lmer5) ## 
#
qqnorm(resid(lmer1))
plot(lmer1) #


# Table 2
lmer1 %>% tbl_regression(intercept=T)  ## manually add LRT values to this


```


```{r, include=F, echo=F}
emmeans(lmer1, "mtype", "mimic")
```
We then focused only on syrphids so that we could use phylogenetic information to ask whether mimics and non-mimics had diverged in preference, and whether body size as well as morphotype could explain flower colour preference. 

Phylogenetic models of yellowness preference were practically indistinguishable from each other according to their DIC, including the model with no explanatory factors at all (delta DIC < 2.5).  Across the phylogeny, syrphids of all morphotypes, mimics and non-mimics, preferred similar flower yellowness (see table 4). In the model with technically the lowest DIC (delta ~ 0.5), larger syrphids had a very slight tendency to prefer less yellow flowers which we regard as negligible.

Table 3. Table of fitted phylogenetic models of yellowness preference among syrphids, showing coefficients and ranked DIC values.

```{r mcmc model for absolute PCA1 value among syrphids including non mimics, include=F , echo=F}
  
## weakly informative prior
prior	<-list(G=list(G1=list(V=1, nu=0.2)), R=list(V=1, nu=0.2))  

# Run MCMC models
m1.raw.pca1	<-	MCMCglmm(pca1 ~ group * meanhw, random=~animal, family="gaussian", pedigree=tree.syrph, nodes='TIPS',data=dat.syrph, prior=prior, verbose=F, thin=50, nitt=100000, burnin=2000)
m2.raw.pca1	<-	MCMCglmm(pca1 ~ group + meanhw, random=~animal, family="gaussian", pedigree=tree.syrph, nodes='TIPS',data=dat.syrph, prior=prior, verbose=F, thin=50, nitt=100000, burnin=2000)
m3.raw.pca1	<-	MCMCglmm(pca1 ~ group         , random=~animal, family="gaussian", pedigree=tree.syrph, nodes='TIPS',data=dat.syrph, prior=prior, verbose=F, thin=50, nitt=100000, burnin=2000)
m4.raw.pca1	<-	MCMCglmm(pca1 ~ meanhw        , random=~animal, family="gaussian", pedigree=tree.syrph, nodes='TIPS',data=dat.syrph, prior=prior, verbose=F, thin=50, nitt=100000, burnin=2000)
m5.raw.pca1	<-	MCMCglmm(pca1 ~ 1        , random=~animal, family="gaussian", pedigree=tree.syrph, nodes='TIPS',data=dat.syrph, prior=prior, verbose=F, thin=50, nitt=100000, burnin=2000)

## DIC comparison
DIC(m1.raw.pca1, m2.raw.pca1, m3.raw.pca1, m4.raw.pca1, m5.raw.pca1)

## All are similar but technically best model is m4.raw.pca1
summary(m4.raw.pca1)

## Table 3
model.sel(m1.raw.pca1, m2.raw.pca1, m3.raw.pca1, m4.raw.pca1, m5.raw.pca1, rank="DIC") %>% 
  mutate_if(is.numeric, round, 2) %>% 
  gt()


```

Figure 3: Yellowness preferences by morphotype and mimicry status. Syrphids (mimetic and non-mimetic) showed similar yellowness preference, while hymenopteran models differed (see text for details).

```{r figure:PCA1 raw values by model/mimic/nonmimic, include=T, echo=F}
fig3_base <- ggplot(
        data=dat %>% 
            mutate(mimic=recode(mimic, 
                          "mim" = "Mimic", 
                          "mod" = "Model",
                          "non" = "Non-mimic")), 
        aes(y=pca1, x=mtype, fill=mimic))

fig3 <-
fig3_base + geom_boxplot(outlier.shape = NA, notch=T)  +
          scale_x_discrete(labels=c('Bumblebee','Honeybee','Solitary bee','Social wasp','Non-mimic\nhoverfly')) +
          xlab('Morphology') + 
          ylab('PCA1 (Yellowness)') +
          theme_classic() +
          theme(legend.title=element_blank())

fig3
```

There were significant differences among groups in blueness preference (Linear Mixed model, X2=30.7, df=8, p<0.001). Bumblebee models and mimics preferred flowers with more blue than others. Honeybee and social wasp morphotypes were intermediate. Solitary bee morphotypes and non-mimic hoverflies preferred flowers with the least blue. 

When we excluded non-mimics and looked specifically at effects of mimicry and morphotype, there was no interaction between "model/mimic" and "morphotype", and also no effect of "model/mimic". This indicated that models tended to choose flowers with similar B values to those chosen by mimics.  In contrast, there was a strong effect of morphotype (X2=26.0, df=3, p<0.0001), showing that different hymenopteran groups (and their syrphid mimics which follow their preferences) tend to choose flowers with distinctive blueness values.

Table 4. Linear mixed-effect model of flower blueness against morphotype, with species as random effect.
```{r, include=F , echo=F}
lmer1 <- lmer(pca2 ~ group + (1 | species),  data=dat) 
lmer2 <- lmer(pca2 ~ 1 + (1 | species),  data=dat) 

anova(lmer1, lmer2) # 
#lmer2: pca2 ~ 1 + (1 | species)
#lmer1: pca2 ~ group + (1 | species)
#      npar    AIC    BIC  logLik deviance  Chisq Df Pr(>Chisq)    
#lmer2    3 2516.7 2531.1 -1255.3   2510.7                         
#lmer1   11 2501.9 2554.8 -1240.0   2479.9 30.721  8  0.0001575 ***

emmeans(lmer1, "group")

# group     emmean    SE   df lower.CL upper.CL
# bbeemim  -0.3946 0.136 75.7  -0.6646  -0.1247
# bbeemod  -0.4202 0.136 75.7  -0.6902  -0.1503
# hbeemim  -0.1875 0.136 75.7  -0.4574   0.0825
# hbeemod   0.1160 0.180 53.4  -0.2456   0.4776
# non       0.1629 0.168 44.0  -0.1747   0.5005
# sbeemim   0.2062 0.136 75.7  -0.0638   0.4761
# sbeemod   0.4104 0.135 78.8   0.1425   0.6783
# swaspmim  0.0520 0.136 75.7  -0.2180   0.3219
# swaspmod -0.0534 0.136 75.7  -0.3234   0.2165

# Table 4
lmer1 %>% tbl_regression(intercept=T)  # Manually add LRT values to this from anova() above
```
Table 5. Linear mixed-effect model of flower blueness against morphotype and "model or mimic", with model-mimic pair as random effect.
```{r lmer analysis - PCA2 (blue) ~ "model group" * "model/mimic" removing nonmimics, include=F , echo=F}
lmer1b <- lmer(pca2 ~ mtype * mimic + (1 | pair_species), data=dat.mimics.models) 
lmer2b <- lmer(pca2 ~ mtype + mimic + (1 | pair_species), data=dat.mimics.models)
lmer3b <- lmer(pca2 ~         mimic + (1 | pair_species), data=dat.mimics.models)
lmer4b <- lmer(pca2 ~ mtype         + (1 | pair_species), data=dat.mimics.models)

anova(lmer1b, lmer2b) ## NS
anova(lmer2b, lmer3b) ## Effect of who you look like
anova(lmer2b, lmer4b) ## No Effect of "model or mimic"
#
qqnorm(resid(lmer1b))
plot(lmer1b) # 

# Table 4
lmer2b %>% tbl_regression(intercept=T)   # Manually add LRT values to this from anova() above
  
```

```{r, include=F , echo=F}
emmeans(lmer1b, "mtype")
```

The three phylogenetic models of blueness preference containing "morphotype" were indistinguishable from each other by their DIC ("morphotype only", "morphotype + body size", "morphotype X body size"; delta < 0.57).  However, all were better than models without "morphotype" by > 4 DIC points (Table 6). This indicated that (as with the non-phylogenetic models above) morphotype, but not body size, was associated with blueness preference. Bumblebee mimics and, marginally, honeybee mimics had different blueness preferences from those of non-mimics, and from those of mimics of solitary bees and social wasps (see MCMCglmm table 7).

Table 6. Table of fitted phylogenetic models of blueness preference among syrphids, showing model terms and ranked DIC values. A + sign indicates presence of a term in a model.

Table 7. Table of fixed coefficients for top phylogenetic model of blueness preference among syrphids

```{r mcmc model for absolute PCA2 value among syrphids including non mimics, include=F , echo=F}
  
phen.var<-var(dat.syrph$pca2,na.rm=TRUE)

## weakly informative prior
prior	<-list(G=list(G1=list(V=1, nu=0.2)), R=list(V=1, nu=0.2))  

dat.syrph$group <- relevel(dat.syrph$group, ref = 'non')

# Run MCMC models
m1.raw.pca2	<-	MCMCglmm(pca2 ~ group * meanhw, random=~animal, family="gaussian", pedigree=tree.syrph, nodes='TIPS',data=dat.syrph, prior=prior, verbose=F, thin=50, nitt=100000, burnin=2000)
m2.raw.pca2	<-	MCMCglmm(pca2 ~ group + meanhw, random=~animal, family="gaussian", pedigree=tree.syrph, nodes='TIPS',data=dat.syrph, prior=prior, verbose=F, thin=50, nitt=100000, burnin=2000)
m3.raw.pca2	<-	MCMCglmm(pca2 ~ group         , random=~animal, family="gaussian", pedigree=tree.syrph, nodes='TIPS',data=dat.syrph, prior=prior, verbose=F, thin=50, nitt=100000, burnin=2000)
m4.raw.pca2	<-	MCMCglmm(pca2 ~ meanhw        , random=~animal, family="gaussian", pedigree=tree.syrph, nodes='TIPS',data=dat.syrph, prior=prior, verbose=F, thin=50, nitt=100000, burnin=2000)
m5.raw.pca2	<-	MCMCglmm(pca2 ~ 1        , random=~animal, family="gaussian", pedigree=tree.syrph, nodes='TIPS',data=dat.syrph, prior=prior, verbose=F, thin=50, nitt=100000, burnin=2000)

DIC(m1.raw.pca2, m2.raw.pca2, m3.raw.pca2, m4.raw.pca2, m5.raw.pca2)

summary(m3.raw.pca2)

## Table 6
model.sel(m1.raw.pca2, m2.raw.pca2, m3.raw.pca2, m4.raw.pca2, m5.raw.pca2, rank="DIC") %>% 
  mutate_if(is.numeric, round, 2) %>% 
  gt()


## Table 7
summary(m3.raw.pca2)$solutions %>% 
  as.data.frame %>% 
  mutate_if(is.numeric, round, 2) %>% 
  gt
```

Figure 4. Blueness preferences by morphotype and mimicry status. Syrphid mimics were different from non-mimics, and tended to follow the preferences of their models such that morphotypes differed but models and mimics were statistically similar (see text for details).

```{r figure:PCA2 raw values by model/mimic/nonmimic, include=T, echo=F}
fig4_base <- ggplot(data=dat %>% 
            mutate(mimic=recode(mimic, 
                          "mim" = "Mimic", 
                          "mod" = "Model",
                          "non" = "Non-mimic")), 
        aes(y=pca2, x=mtype, fill=mimic))


fig4 <-
fig4_base + geom_boxplot(outlier.shape = NA, notch=T)  +
          scale_x_discrete(labels=c('Bumblebee','Honeybee','Solitary bee','Social wasp','Non-mimic\nhoverfly')) +
          xlab('Morphology') + 
          ylab('PCA2 (Blueness)') +
          theme_classic() +
          theme(legend.title=element_blank())

fig4

```
````
