## Supplementary material for "Internet images reveal bumblebee-mimicking hoverflies have evolved to follow flower colour preferences of their models": Readme.txt

# README

Title of study
Internet images reveal bumblebee-mimicking hoverflies have evolved to follow flower colour preferences of their models.

Short summary of the study
We gathered internet photographs of flower-visiting bees and wasps, their hoverfly mimics, and non-mimicking hoverflies, and compared the colours of the visited flowers. We predicted that, driven by selection for enhanced mimicry, bee-mimicking hoverflies would have switched flower preference from yellow to blue, to follow the flower choices of their models.

Files included 
## Description/Comments on each file

2024 Syrphid behavioural mimicry - results and code.Rmd 
## R Markdown file that reproduces the results section of the paper from the files enclosed here.
## Some figures are produced in R but compiled manually - this is indicated in the comments.

Table S1 model-mimic ids.xls 
## Table of mimics and corresponding models compiled from literature. 
## References are given in the separate worksheet.
## File is not used in the code so there is no corresponding .csv file

Table S2 head width.xlsx 
Table S2 head width.csv 
## Table of head widths for syrphids in dataset
## These data are a combination of direct measurement and interpolation based on other traits (see text for details)
#	species				Species of Syrphid
#	head width (mm)			Head width in mm
#	Interpolated of measured	Whether interpolated or measured (see text for details)


Table S3 main data.xlsx
Table S3 main data.csv
## Main data table - data on morphotype group (see text for details), model species (for mimics), and median flower colour for each photograph.
## Metadata are in a separate worksheet in the .xlsx file and are replicated here:
#	Field		Description
#	species		Species name of mimic or model
#	group		morphotype group (see text for details): bbee = bumblebee; hbee = honeybee; sbee = solitary bee; swasp = social wasp; mim = mimic; non = non-mimic; mod = model
#	pair_species	Model species (see text and Table S1 for details)
#	pair2		Model search string; for use in grouping model-mimic pairs (see code for details)
#	r		Median red content of flower cutout (0-255)
#	g		Median green content of flower cutout (0-255)
#	b		Median blue content of flower cutout (0-255)

Supplementary file S4 - phylogeny.nex
## Syrphid phylogeny at the genus level in NEXUS format
## For analysis, species in the dataset are added in to this at the coding stage (see code for details).
